# PBRM1-Dependent PBAF Targeting is Required for EMT and Metastasis in Breast Cancer

**DOI:** 10.1101/2025.10.19.683137

**Authors:** Alisha Dhiman, Mitchell G. Ayers, Jamie L. McCuiston, Aparna B. Shinde, Saeed Salehin Akhand, Guanming Jiao, Juan Jauregui-Lozano, Marco Hadisurya, Elizabeth G. Porter, W. Andy Tao, Vikki M. Weake, Yucheng Zhang, Sagar M. Utturkar, Matthew R. Maraunde, Michael K. Wendt, Emily C. Dykhuizen

## Abstract

SWI/SNF chromatin remodelers utilize ATP to mobilize nucleosomes on DNA and are represented by three biochemically distinct subcomplexes, the more abundant cBAF and the less abundant PBAF and GBAF subcomplexes. Patient mutations and genetic studies have identified important roles for PBAF subunits in development and disease; however, relating PBAF-mediated phenotypes to biochemical function in chromatin regulation and gene expression has been challenging. Further complicating matters, cell-based systems often do not reflect the phenotypes and genotypes observed with PBAF mutations *in vivo*. Here we show that the PBRM1 subunit of PBAF is critical for the completion of TGFB1-mediated epithelial-mesenchymal transition of mammary cells in vitro as well as the metastasis of murine breast cancers in vivo. Using epigenomics to profile different stages of EMT, we find that PBRM1 is necessary for targeting PBAF to inducible promoters marked by H3K14ac alone. We further find that PBRM1 facilitates DNA accessibility at sites bound by TGFβ1-inducible transcription factors, such as Atf3, for the induction of genes involved in migration, cell survival, and inflammation. Our model allows us to separate constitutive vs inducible gene expression to help explain some of the context-dependent phenotypes observed with PBRM1 deletion. In addition, we provide evidence that while PBRM1 deletions can promote the initiation of certain cancers in early stages, PBAF may be a vulnerability in late-stage metastatic cancers.

## Introduction

Mammalian SWI/SNF complexes (also termed BAF for Brg1/Brm associated Factors) are multi-subunit complexes that use the energy of ATP to mobilize nucleosomes and create regions of DNA accessibility. SWI/SNF complexes are highly heterogeneous, with hundreds of different combinations of subunits that facilitate cell-type specific transcription programs^1^. In addition, all cells contain three biochemically distinct SWI/SNF complexes that can be differentiated by size and abundance: cBAF, PBAF, and most recently, GBAF^2^ (also called ncBAF^3,4^). cBAF contains the unique subunits ARID1A/B and DPF1/2/3 and is typically the most heterogeneous and abundant of the three SWI/SNF complexes^4^. It is also primarily responsible for the largest phenotypic and transcriptional effects in the cell^5–7^. The majority of SWI/SNF-driven chromatin accessibility regulation is mediated through cBAF-driven remodeling at enhancers^8–13^. In contrast, PBAF, with unique subunits PHF10/ARID2/BRD7/PBRM1, and GBAF, with unique subunits GLTSCR1or1L/BRD9, are dispensable for viability in most cells and seem to contribute very little to overall accessibility^5,14^. For GBAF, this is likely due to low expression and the lack of SMARCB1, which is critical for nucleosome eviction^15,16^. PBAF, however, has critical interactions between core subunits^16,17^ and the nucleosome^17,18^, and cBAF-comparable remodeling activity *in vitro*^19,20^. The reasons behind PBAF’s relatively minor contribution to accessibility in cells remain poorly understood.

Most functional studies of PBAF focus on the roles of PBRM1 and ARID2 in cancers where they are highly mutated^21,22^. In these cancer backgrounds, the reported effects of PBAF subunit depletion are variable and context-dependent, often failing to display obvious tumor suppressive phenotypes^23,24^. In cell systems where PBAF does have a tumor suppressive phenotype, the effects are often gain-of-function, whereby PBAF subunit deletion leads to an increase in the activity of other transcriptional factors. For instance, HIF1 or NFκB activity is increased in renal cancers with PBRM1 deletion^25,26^ and cBAF/AP-1 activity is increased in melanoma with ARID2 deletion^27^. This has led to the hypothesis that PBAF primarily acts as a transcriptional repressor^28,29^ by moving nucleosomes to block transcription at sites of DNA damage^28,29^ or to facilitate binding of repressive transcription factors, such as REST^14^. Mechanistic insight into how PBAF represses transcription in other systems is limited. Many of these studies provide valuable support that deletion of PBAF facilitates the activation of oncogenic pathways in specific cancer contexts^25–28^; however, they don’t adequately address how these phenotypes relate to the biochemical mechanism of PBAF-mediated chromatin binding and remodeling.

In our previous publication, we found that deletion of PBRM1 in epithelial cells increases proliferation under nutrient replete growth conditions but decreases viability of the same cell type upon induction of cell stress ^24^. Here we report that these same epithelial cells with PBRM1 deletion are unable to survive TGFβ1 treatment and complete epithelial-mesenchymal transition (EMT). We use a multi-omics approach to define the transcriptional role for PBRM1 in gene activation during EMT and the related nucleosome binding and chromatin remodeling changes required for this cell state transition. We further demonstrate that this function is essential for metastasis in a variety of mouse models, implicating PBAF as a potential target in metastatic disease.

## Materials and Methods

RNAseq data has been submitted to GEO under GSEXXXXXXX, ATACseq data under GSEXXXXXXX, ChIPseq data will be submitted to GEO once government reopens

### Cell lines and culture conditions

NMuMG cells were purchased from ATCC and grown in DMEM (Corning) supplemented with 10% fetal bovine serum (Corning), 10 µg/mL Insulin (Sigma), 1% L-glutamine (Corning), 1% antibiotics (100 units/mL penicillin and 100 g/mL streptomycin; Corning), and 1% Sodium Pyruvate (Corning)

The EGFR transformed NME cell line with stable luciferase expression has been reported before^30^ and was cultured similar to NMuMG cells as described above.

4T1 and 4T07 cells stably expressing luciferase^31,32^ were grown in DMEM (Corning) supplemented with 10% fetal bovine serum (Corning), 1% L-glutamine (Corning), 1% antibiotics (100 units/mL penicillin and 100 g/mL streptomycin; Corning), and 1% Sodium Pyruvate (Corning).

Renca cells were grown in RPMI-1640 (Corning) supplemented with 10% fetal bovine serum (Corning), 1% Non-essential amino acids (Corning), 1% L-glutamine (Corning), 1% antibiotics (100 units/mL penicillin and 100 g/mL streptomycin; Corning), and 1% Sodium Pyruvate (Corning).

MCF10A-CA1h, MCF10A-CA1a, and HEK293T were cultured in DMEM (Corning) supplemented with 10% fetal bovine serum (Corning) 1% L-glutamine (Corning), 1% antibiotics (100 units/mL penicillin and 100 g/mL streptomycin; Corning), and 1% Sodium Pyruvate (Corning).

All cell lines were grown at 37 °C in a humidified atmosphere in a 5% CO2 incubator. All the media were supplemented with 1:10,000 dilution of Plasmocin™ (InvivoGen), and routinely checked for mycoplasma contamination.

### Generation of cell lines

Constitutive knockdown was performed using shRNA-mediated knockdown with lentiviral construct pLKO.1. Doxycycline-inducible knockdown was performed using the lentiviral construct pLKO-tet-ON^33^. The shRNA constructs contain the following mature antisense sequences:

Mouse Pbrm1: (TRCN0000081820) TTCTAGGTTGTATGCCTGTCG

Mouse Brd7: (TRCN0000030015) ATAATCATGGAGTAGCCAGGC

Mouse Arid1A: (TRCN0000071395) ATTGTAGGTCATGTCATTTCG

Mouse Atf3: (TRCN0000082131) TTGTTTCGACACTTGGCAGCA

Scramble control hairpin: GCTACACTATCGAGCAATT

NMuMG Pbrm1 KO cell line generation has been described before^24^. The same constructs and methodology were used for generating Renca, 4T1 and NME Pbrm1 KO cell lines.

Renca parental cells were first transduced with lentiviral particles for the dual reporter construct pFU-Luc2-eGFP (L2G)^34^ (a kind gift from Huiping Lui) and the GFP expressing cells were selected using FACS. These GFP positive cells were then transduced with either shRNA lentiviral particles or used for generating Pbrm1 KO cells.

Atf3 re-expression was done by transient transfection using the addgene construct# 26115 (pRK-ATF3 was a gift from Yihong Ye)^35^, stable clones were selected by continuous G418 administration (600 µg/mL) and used for experiments.

After performing any genetic perturbation, cell line growth rates were compared by seeding a fixed number of cells on day 0 and then subsequent cell counting using trypan blue at the indicated time-points. For longer time frames, a proportion of cells was re-seeded at the time of counting to take to the next time point. Two-way ANOVA was done for statistical comparison.

### Lentiviral Infection

HEK293T cells were transfected with knockdown lentivirus constructs along with packaging vectors (pMD2.G and psPAX2 for pLKO.1; pCMV-8.2ΔVPR and pCMV-VSV-G for pLKO-tet-ON). After 48 h, the supernatant was collected and concentrated by ultracentrifugation (17,500 rpm with Beckman rotor SW 32 Ti/∼52,000 x g and resuspended in 200 µL of PBS. Cells were transduced with concentrated virus using spinfection (1500 rpm/∼250 x g in swing bucket centrifuge for 1 h) and incubated for 48 h. Cells were selected with puromycin (2 µg/mL) (Sigma-Aldrich) and hygromycin (200 µg/mL) (Corning) where applicable. The efficiency of all constructs was confirmed by immunoblotting.

### Immunoblotting

Cells were given the requisite treatments for the indicated time periods, followed by harvesting by trypsinization. Whole cell extracts were prepared by dissolving the cell pellets in RIPA buffer (50 mM Tris (pH 8.0), 150 mM NaCl, 0.1% SDS, 0.5% Na Deoxycholate, 1% NP-40) supplemented with freshly added PMSF, aprotinin, leupeptin and pepstatin, and incubation for 30 min at 4 °C. Nuclear lysates were prepared by resuspending cells in Buffer A (20 mM HEPES (pH 7.9), 25 mM KCl, 10 % glycerol, 0.1% Nonidet P-40 with PMSF, aprotinin, leupeptin, and pepstatin) at a concentration of 20 million cells/mL. The cells were kept on ice for 5 min, and nuclei were isolated by centrifugation at 600 x g for 10 min. The nuclei pellet was resuspended in chromatin IP buffer (20 mM HEPES (pH 7.9), 150 mM NaCl, 1% Triton X-100, 7.5 mM MgCl_2_, 0.1 mM CaCl_2_ with PMSF, aprotinin, leupeptin, and pepstatin). The lysates were centrifuged at 21,000 g for 30 min at 4 °C and the supernatants were preserved. Protein concentration estimations for the supernatants were done using BCA protein assay kit (Pierce Biotechnology) with BSA as the standard and equal amounts of proteins were mixed with 4× lithium dodecyl sulfate sample buffer containing 10 % 2-merchaptoethanol. The proteins were denatured for 5 min at 95 °C, separated on a 4-12 % bis-tris gradient protein gel and transferred to PVDF membrane. The membrane was blocked with 5 % BSA in TBS containing 0.1% Tween 20 for 30 min at room temperature and then incubated with primary antibodies overnight at 4 °C. The primary antibodies were detected by incubating the membranes in goat anti-rabbit or goat anti-mouse secondary antibodies (LI-COR Biotechnology, Lincoln, NE) conjugated to IRDye 800CW or IRDye 680RD, respectively, for 1 h at room temperature, and the signals were visualized using Odyssey Clx imager (LI-COR Biotechnology).

### Immunoprecipitation (IP)

Cell treatments and nuclear lysates were prepared in chromatin IP buffer as described above and treated with 4U/mL Turbo DNase (Ambion) for 30 min at 4 °C. The extracts were cleared by centrifugation 21,000 x g at °4 C for 30 min. Protein estimation was done on the clarified supernatants using BCA protein assay kit (Pierce Biotechnology) with BSA as the standard. For ATF3 IP, 5% input was taken out and 500 µg nuclear lysate/sample was used for setting up IP using ATF3 antibody (ab207434 Abcam) or Normal IgG (2729S CST) and rotated overnight at 4 °C. Equilibrated protein A beads (20 µL/IP) were added to the lysates and rotated at 4 °C for another 2 h. The bead-bound complexes were washed twice with chromatin IP buffer and thrice with high stringency buffer (20 mm HEPES (pH 7.9), 500 mM NaCl, 1% Triton X-100, 0.5% sodium deoxycholate, 1 mM EDTA with PMSF, aprotinin, leupeptin, and pepstatin) followed by boiling in 1X lithium dodecyl sulfate loading dye at 80 °C for 10 min and immunoblotting. For PHF10 IP-MS, a 2% input was removed and 500 µg lysate/sample was used in an IP with PHF10 antibody (PA5-30678 Invitrogen) and rotated overnight at 4 °C. Equilibrated protein A beads (15 µL/IP) were added to the lysates and rotated at 4 °C for another 2 h. The bead-bound complexes were washed thrice with chromatin IP buffer followed by three 1X PBS washes and submitted for IP-MS.

### LC-MS Sample Preparation

The phase-transfer surfactant (PTS) buffer containing 12 mM sodium deoxycholate, 12 mM sodium lauroyl sarcosinate, 10 mM TCEP, and 40 mM chloroacetamide (CAA) in 50 mM Tris·HCl, pH 8.5 was added to the magnetic beads. The proteins were incubated for 10 min at 95°C and digested on beads with Lys-C (Wako) at 1:100 (wt/wt) enzyme-to-protein ratio for 5 h at 37 °C. Furthermore, trypsin was added to a final 1:50 (wt/wt) enzyme-to-protein ratio for overnight digestion at 37 °C. The supernatant containing the digested proteins was separated from the magnetic beads with the help of a magnetic rack separator and processed further. Then, a final concentration of 1% trifluoroacetic acid (TFA) was added to acidify the samples. An ethyl acetate solution was added at a 1:1 ratio to the samples. The solution was vortexed for 2 min and then centrifuged at 20,000 x g for 3 min to separate the aqueous and organic phases. The top layer (the organic phases) was removed, and the aqueous phase was collected, dried down in a vacuum centrifuge, and desalted using TopTip C18 tips (Glygen) according to the manufacturer’s instructions. The desalted samples were dried completely in a vacuum centrifuge.

### LC-MS Analysis

The proteomics samples were spiked with an 11-peptide Retention Time internal standard (Biognosys) to normalize the LC-MS signal between sample runs. All samples were loaded into an Easy-nLC 1000 (Thermo Fisher Scientific) and separated with a 45-cm packed column (360-µm o.d. × 75-µm i.d.) containing C18 resin (2.2 µm, 100Å; Michrom Bioresources) with a heated 30-cm column heater (Analytical Sales and Services) set to 50°C. The mobile phase buffer contained 0.1% formic acid in HPLC grade water (Buffer A) with an eluting buffer containing 0.1% formic acid in 80% (vol/vol) acetonitrile (Buffer B) run with a linear 60-min gradient of 10-35% Buffer B at a flow rate of 300 nL/min. The HPLC was coupled with an LTQ-Orbitrap Velos Pro mass spectrometry (Thermo Fisher Scientific). The mass spectrometry was run in data-dependent mode with a full-scan MS (from m/z 350 to 1,500 with a resolution of 30,000), followed by MS/MS of the 10 most intense ions subjected to collision-induced dissociation (CID) fragmentation. CID fragmentation was performed and acquired in the linear ion trap (minimal signal threshold 1,000 counts, normalized collision energy (NCE) 30%, activation Q 0.25, activation time 10 ms, default charge state 3, isolation window 3 m/z, and dynamic exclusion 60 s).

### LC-MS Data Processing

The raw files were searched against the mouse Swiss-Prot database with no redundant entries, using Sequest and Byonic (Protein Metrics) search engines loaded into Proteome Discoverer 2.3 software (Thermo Fisher Scientific). MS1 precursor mass tolerance was set at 10 ppm, and MS2 fragment tolerance was set at 0.6 Da. In the processing workflow, search parameters for both search engines were performed with full trypsin/P digestion, a maximum of two missed cleavages allowed, a static modification of carbamidomethylation on cysteines (+57.0214 Da), and variable modifications of oxidation (+15.9949 Da) on methionine residues and acetylation (+42.011 Da) at N-terminus of proteins. The false-discovery rates (FDR) of peptide spectrum matches (PSMs), peptides, and proteins were set at 0.01 (strict) and 0.05 (relaxed). All protein and peptide identifications were grouped, and any redundant entries were removed. Unique peptides and unique master proteins were reported. Finally, the proteomic abundance results were further normalized using the spiked 11-peptide Retention Time internal standard.

### TGFβ1 treatment assays

NMuMG cells were seeded in 6-well plates with the indicated concentrations of TGFβ1 (R&D systems, Catalog # 7754-BH) to induce EMT and cultured for 2-10 days as detailed in the respective figure legends. To assess if the EMT was reversible, the cells were first treated with TGFβ1 for 5 days and then TGFβ1 was removed while culturing the cells in the same way as described above. The cells were trypsinized and counted using the trypan blue method every 48-72h and re-seeded in 6-well plates at the same cell number as day 1 at each passage. Cell survival was calculated as % cell number relative to untreated control within the group. Each experiment was done in at least 3 biological replicates. Error bars represent SD, stats were done using multiple unpaired t-tests with Holm-Sidak correction.

### Cell cycle analysis

NMuMG cells were treated with TGFβ1 for 0-72h as described above and analyzed using the 488 EdU Click Proliferation Kit (BD Biosciences, # 565455) at the indicated time-points. For each time-point, 10 µM Edu was added to the media 1h before harvesting and the cells were processed and frozen until ready for use according to the manufacturer’s instructions. Propidium Iodide/RNase solution (Invitrogen, # F10797) was used for DNA staining. The cells were run on the BD Fortessa LSR flow cytometry cell analyzer and the data were analyzed using FlowJo software. The PI intensity was used to designate G0/G1 and G2/M populations. Each experiment was done in 3 biological replicates. Error bars represent SD, stats were done using multiple unpaired t-tests with Holm-Sidak correction.

### Apoptosis analysis

NMuMG cells were treated with TGFβ1 for 0-5 days as described above and harvested using Accutase (Corning, #25-058-CI) at the indicated time-points. The cells (including floating cells) were processed and stained using the FITC Annexin V Apoptosis Detection Kit I (BD Biosciences, # 556547) according to the manufacturer’s instructions. The cells were immediately run on the BD Fortessa LSR flow cytometry cell analyzer and the data were analyzed using FlowJo software. Each experiment was done in 2 biological replicates. Error bars represent SD, stats were done using multiple unpaired t-tests with Holm-Sidak correction. A portion of the harvested cells above were used for C-PARP immunostaining as an additional assay for apoptosis.

### Invasion assays

NMuMG and 4T1 cells were either left untreated or treated with TGFβ1 for 5 days, before seeding on the transwells for invasion assays. The assay was done using regular Pathclear BME or reduced growth factor Pathclear BME (R&D Systems) diluted to 10% in serum free media, TGFβ1 in serum-free media was added to the top chamber only, 5-10% serum was added to the bottom chamber only and serum free media in both chambers was used as the baseline invasion control. The invading cells were fixed and stained with crystal violet after 48h incubation. The stained cells were photographed, and the stain was redissolved in 0.2% SDS and quantitated by absorbance readings at 570nm. The experiments were done once with 3 technical replicates. Error bars represent SD, stats were done using multiple unpaired t-tests with Holm-Sidak correction.

### In vivo studies

Luciferase expressing 4T1 cells (4T1-L4)^32^ transduced with either pLKO.1, *shPbrm1* (10 mice/group), *shBrd7*, *shArid1A* (5 mice/group) or sgCt, *sgPbrm1*, genetic perturbations were orthotopically engrafted onto the mammary fat pads of 4-6 weeks old female Balb/c mice (Jackson Labs) - 50,000 cells/mouse in 50µL 1X PBS. Primary tumor measurements were done weekly using vernier calipers and the tumors were surgically removed 2 weeks after engraftment. Metastasis development was monitored by weekly bioluminescent imaging using the Advanced Molecular Imager (Spectral Instruments).

4T1-L4 cells expressing either doxycycline-inducible scramble or *shPbrm1* (12 mice/group) genetic perturbations were orthotopically engrafted onto the mammary fat pads of 4-6 weeks old female Balb/c mice (Jackson Labs) - 50,000 cells/mouse in 50 µL 1X PBS. Primary tumor measurements were done weekly using vernier calipers and the tumors were surgically removed after 2 weeks of injections. Each group was then divided into no doxycycline administration (5 mice/group) or doxycycline administration in drinking water (7 mice/group) sub-groups such that average primary tumor size was comparable. Doxycycline water administration (1 mg/mL+5% sucrose in drinking water) was started 24 h after primary tumor removal, with 2 changes/week until the experiment ended. Metastasis development was monitored by weekly bioluminescent imaging using the Advanced Molecular Imager (Spectral Instruments).

Control or Pbrm1-depleted (pLKO.1, *shPbrm1*; 5 mice/group) 4T07 pLuc cells^31^ were injected into the tail veins of female Balb/c mice (1.0 x 10^6^ cells in 100µL 1X PBS) and pulmonary tumor development was monitored by weekly bioluminescent imaging.

Control or Pbrm1-depleted (sgCt, *sgPbrm1*) NME pLuc cells^30^ were orthotopically engrafted onto the mammary fat pads of 15-16 week old female NRG mice (Purdue Biological Evaluation core, 8 mice/group)-1.1 X 10^6^ cells/mouse in 50 µL 1X PBS. Primary tumor measurements were done weekly using vernier calipers and the tumors were surgically removed after 4 weeks of injections. Metastasis development was monitored by weekly bioluminescent imaging. At the time of necropsy, a portion of the lungs was saved for *ex vivo* sub-culture with antibiotic selection to enrich tumor derived cells and immunoblotting was performed for Pbrm1 protein expression.

Control or Pbrm1-depleted (pLKO.1, *shPbrm1*; 6 mice/group) and control or Pbrm1-deleted (sgCt, *sgPbrm1*; 7 mice/group) Renca FF-GFP-pLuc cells were injected into the lateral tail vein of male Balb/c mice (100,000 cells/mouse) in 100µL 1X PBS and pulmonary tumor development was monitored by weekly bioluminescent imaging. For the experiment with control and Pbrm1-deleted Renca cells, at the time of necropsy, a portion of the lungs was saved for ex-vivo subculture with antibiotic selection to enrich tumor derived cells and immunoblotting was performed for Pbrm1 protein expression.

Control or Pbrm1-depleted (pLKO.1, *shPbrm1*) Renca pLuc cells were injected into the lateral tail vein of male nu/nu mice (5 mice/group (Taconic); 100,000 cells/mouse) in 100µL 1X PBS and metastasis development was monitored by weekly bioluminescent imaging.

Upon necropsy, primary tumors (if applicable) and lungs from all animals were removed, fixed in 10% formalin for 24h and dehydrated in 70% ethanol for visualization of pulmonary metastatic nodules and further histologic analyses. All animal studies were performed in accordance with the animal protocol procedures approved by the Institutional Animal Care and Use Committee of Purdue University. Statistical analyses were performed using Welch’s t-test and two-way ANOVA with multiple comparisons as indicated in the figure legends

### qRT-PCR

RNA was extracted using TRIzol (Ambion, Inc.). cDNA was synthesized using Verso cDNA synthesis kit (Thermo Scientific) using 3:1 mix of random hexamers and oligo dT primers as per manufacturer’s recommendation. Specific targets were amplified using PowerUp SYBR Green Master Mix (Applied Biosystems) in Bio-Rad CFX qPCR instrument. The following qPCR primers were used: mAtf3 FP: GAGGATTTTGCTAACCTGACACC; mAtf3 RP: TTGACGGTAACTGACTCCAGC; mOaz1 FP: GTGGTGGCCTCTACATCGAG; mOaz1 RP: GTTGCTCCTCTGCGAACTCTA.

### RNA-seq

NMuMG cells were seeded in 6-well plates with or without TGFβ1 as described above. Cells were harvested at 24, 48 and 72 h post-treatment in trizol and total RNA was isolated using the PureLink RNA isolation kit (Invitrogen, #12183018A). Downstream library construction, sequencing on Novaseq 6000 and data analysis was done by Novogene. 4T1 cells were seeded with or without TGFβ1 for 4 days and processed as above for total RNA isolation. Library construction was done using Illumina Stranded mRNA prep, ligation kit (Illumina, #20040534) according to manufacturer’s instructions and checked for quality using Qubit and Agilent Bioanalyzer by Purdue Genomics Core Facility. Libraries were sequenced using 150bp PE sequencing on NovaSeq 6000 platform (Novogene, Sacramento, CA). All experiments were performed in three biological replicates.

### RNA-seq analysis

NMuMG *sgPbrm1* and sh*Atf3* datasets were performed by Novogene: Raw reads were trimmed for adapter, poly-N and low quality sequences using fastp, and mapped to mouse mm10 genome build using STAR^36^ (v2.6.1d) software (NMuMG *sgPbrm1* datasets, mismatch=2) or Hisat2^37^ (v2.0.5) software (NMuMG shATF3 datasets, default parameters). Differential expression analysis was done using DESeq2^38^ R package (v1.20.0), and the resulting *p-*values were adjusted using the Benjamini and Hochberg’s approach for controlling the False Discovery Rate (FDR). Heatmaps were visualized using Heatmap2^39^ and Venn diagrams were visualized using Eulerr^40^.

For 4T1 sh*Pbrm1* datasets: Raw reads were trimmed for adapter sequences and low-quality bases using trim-galore^41^ (v0.6.7) and mapped to mouse mm10 genome build using STAR^36^ (v2.7.10a). Reads with MAPQ >10 were used for downstream analysis. FeatureCounts from subread^42^ (v2.0.1) was used to map reads to corresponding genes according to GENECODE Mus_musculus.GRCm38.102 annotation. Differential gene expression analysis was performed by edgeR^43^ (v3.38.0). Read counts were normalized using the trimmed mean of M-values (TMM) method, after filtering out lowly expressed genes with the filterByExpr function using its default settings. Genes with FDR-adjusted *p*-values < 0.01 and absolute log_2_ fold changes > 1 were considered as DEGs.

### ATAC-seq

NMuMG cells were seeded in 6-well plates with or without TGFβ1 as described above. Cells were harvested at 24, 48 and 72h post-treatment and processed according to the Omni-ATAC protocol^44^. The final libraries were subjected to double size selection according to a published protocol^45^ and quality checked using Qubit and Agilent Bioanalyzer by Purdue Genomics Core Facility. Libraries were sequenced using 150bp PE sequencing on NovaSeq 6000 platform (Novogene, Sacramento, CA). All experiments were performed with three biological replicates.

### ATAC-seq data processing

Quality trimming (adapter removal, phred quality cutoff >=Q30, minimum read length 50 bp after quality filtering) of raw reads was performed using fastp^46^ (v0.19.5) tool. Trimmed reads were mapped to mouse mm10 genome build using bowtie2^47^ (v2.3.3) aligner. Peak calling was performed with Genrich in ATAC-seq mode with parameters (-j -r -e chrM -m 30 -v -k -q 0.05) that accounts for ATAC-seq shifting, minimum MAPQ 30, q-value cutoff 0.05 and removal of mitochondrial, duplicate reads, non-unique mapped reads and improperly paired reads. Peaks were annotated using R-package ChIPseeker^48^ (v3.15). Differential peak calling was performed using MACS2^49^ (v2.2.9.1) using default parameters in the galaxy interface^50^. PCA plot was generated using DiffBind^51^ (v3.19.0).

### DiffTF analysis on ATAC-seq

DiffTF^52^ analysis was performed as previously published^53^ in basic mode using the Jan 2012 GRCm38 mm10 target genome and motifs from the JASPAR library^54^ with default parameters (p-value<0.00001, Bg base composition 0.29,0.21,0.21,0.29). Identified TFs were classified as significant if they had an FDR lower than 0.001 in at least one of the pair-wise comparisons between sgCt and sg*Pbrm1*.

### ChIP-seq/ChIP-qPCR

NMuMG cells were seeded in 15cm tissue culture plates with or without TGFβ1 as described above, such that they were ∼80-90% confluent on the day of harvesting (30-40 million cells/15cm at harvest). Cells were either cross-linked with 1% formaldehyde only (histone modifications, Fosb, Fosl2 ChIP-seq) for 10 min at RT, or sequentially dual crosslinked (Brg1, PHF10 and ATF3 ChIP-seq) with 2 mM DSG for 45 min at RT followed by 1% formaldehyde for 10 min at RT as described in ^55^. Crosslinking was quenched with 125 mM Glycine for 10 min at RT. The cells were washed by ice-cold 1 X PBS, scraped from plate, and flash frozen to be stored at −80 °C until ready for processing. Cell pellets were thawed on ice, extracted with nuclear extraction buffer (50mM HEPES-KOH pH 8.0, 140mM NaCl, 1mM EDTA, 10% glycerol, 0.5% NP-40, 0.25% Triton X-100) for 10 min on ice and washed once with nuclear wash buffer (10mM Tris-HCl pH 8.0, 1mM EDTA, 0.5mM EGTA, 200mM NaCl). Nuclear pellets were resuspended in shearing buffer (10mM Tris-HCl pH 8.0, 1mM EDTA, 0.1% SDS) and sonicated using Branson SFX250 at 20% amplitude for 9 mins for dual cross-linked cells and 6 mins for single cross-linked cells at 0.5 s ON and 1.5 s OFF setting to obtain 200-800 bp fragment size. Sonicated chromatin was clarified by high-speed centrifugation and a 50 µL aliquot was reverse cross-linked to check shearing efficiency. The chromatin was quantitated using absorbance at 260 nm and further diluted with 2X ChIP dilution buffer (100 mM HEPES-KOH pH 7.5, 600 mM NaCl, 2 mM EDTA, 2% Triton X-100, 0.2% Sodium Deoxycholate, 0.2% SDS) to obtain equal chromatin amounts in equal volume. Antibodies against BRG1 (ab110641 Abcam), PHF10 (PA5-30678 Invitrogen), ATF3 (ab207434 Abcam), H3K14ac (07-353 Millipore Sigma), H3K18ac (ab1191 Abcam), H3K27ac (ab4729 Abcam), H3K4me (ab8895 Abcam), H3K4me3 (ab8580 Abcam), Fosb (MA5-15056 Invitrogen) and Fosl2 (sc-166102 Santa Cruz) were added. After overnight primary antibody incubation at 4 °C with rotation, Protein A (for rabbit antibodies) or Protein G (for mouse antibodies) Dynabeads were added and mixed using rotation for 2 h at 4 °C. The beads were sequentially washed with low-salt wash buffer (20 mM HEPES-KOH pH 7.5, 0.1% SDS, 0.1% Deoxycholate, 1% Triton, 150 mM NaCl, 1 mM EDTA, 0.5 mM EGTA), high-salt wash buffer (20 mM HEPES-KOH pH 7.5, 0.1% SDS, 0.1% Deoxycholate, 1% Triton, 500 mM NaCl, 1 mM EDTA, 0.5 mM EGTA), LiCl wash buffer (20 mM HEPES-KOH pH 7.5; 0.5% Deoxycholate, 0.5% IGEPAL CA-630; 250 mM LiCl, 1 mM EDTA, 0.5 mM EGTA) and final wash buffer (20 mM HEPES-KOH pH 7.5, 1 mM EDTA, 0.5 mM EGTA). The immunoprecipitated chromatin was eluted once with 200 μL and once with 100 μL elution buffer (100 mM NaHCO3, 1% SDS) for 30 minutes each at 37 °C with shaking. The eluate and the saved input were treated with 2 µL RNase A (10 mg/mL, Thermo Scientific EN0531), 2 µL Proteinase K (20 mg/mL, Thermo Scientific EO0491) and reverse cross-linked at 65 °C for 16 h. DNA was extracted once with phenol:chloroform; once with chloroform and then precipitated by adding 1/10 volume of 3M NaOAc pH 5.2; 1 volume 2-propanol and 2 μL glycogen (20 mg/mL, Thermo Scientific R0561) overnight at −20 °C. After centrifugation at top speed (∼21,000 x g) for 1 h, the pellet was washed with fresh 70% ethanol and then air-dried. DNA was resuspended in low-EDTA TE buffer and DNA quality, and concentration were determined using Qubit and Agilent TapeStation by Purdue Genomics Core Facility. For ChIP-qPCR, 4 µL ChIPed DNA was used per sample. ChIP-seq library preparation was done using Ovation Ultralow System V2 UDI (Tecan Genomics) according to manufacturer’s directions, and the final library was subjected to double sided size selection of 0.65x and 1x. Library preparation was also quality checked and submitted for 150 bp PE sequencing on NovaSeq 6000 platform (Novogene, Sacramento, CA). Primers for ChIP-qPCR: mAtf3 FP: CCTTATCAGGCTGGGAGCCG, mAtf3 RP: GGTGGAGTCATGCCGCTG.

### ChIP-seq data processing

Quality trimming (adapter removal, phred quality cutoff >=Q30, minimum read length 50 bp after quality filtering) of raw reads was performed using fastp^46^ (v0.19.5) tool. Trimmed reads were mapped to mouse mm10 genome build using bowtie2^47^ (v2.3.3) aligner. BAM to BigWig conversion was performed using deeptools^56^ (v3.5.1) with CPM normalization. Peak calling was performed with MACS3^49^ (v3.0.0a7) within Partek Flow (v11.0.24.0325) with -BROAD ON for SWI/SNF subunits. Peaks annotation was performed using R-package ChIPseeker^48^ (v3.15). Peak overlaps were performed using BEDtools^39^ (v2.31.1), heatmaps and metagene analyses were performed using deeptools^56^ (v3.5.4).

### Expression and Purification of human PBRM1 BDs

Constructs encoding codon-optimized ORFs for bacterial expression of human PBRM1 BD2 (addgene #39013), BD3 (addgene #39030), BD4 (addgene #39103), BD5 (addgene #38999) and tandem BD2-5 (SGC construct ID #PB1A-c080) were transformed in BL21(DE3) for BDs, and BL21 Rosetta2 (DE3) pLysS for tandem BD2-5. Protein was induced with 0.1 mM IPTG and expressed at 18 °C overnight. Cultures were resuspended in binding buffer (20 mM Tris pH 8.0, 150 mM NaCl, 25 mM Imidazole, 5% Glycerol with EDTA-free protease inhibitors), passed through a 18½ gauge needle 10-12 times, sonicated at 10% amplitude with 30 sec-ON 30 sec-OFF cycle for a total of 6 min, and centrifuged at 21,000 x g for 1h at 4 °C. An aliquot of the supernatant was removed for the gel, and the rest of the supernatant was incubated with binding buffer-equilibrated Ni^+2^-NTA resin (HisPur Ni-NTA #88221, ThermoFisher Scientific) at 4 °C for 2h with end-to-end rotation. The protein-bound resin was washed three times with wash buffer (20mM Tris pH 8.0, 150mM NaCl, 50mM Imidazole, 5% Glycerol with EDTA-free protease inhibitors), and eluted four times with elution buffer (20mM Tris pH 8.0, 150mM NaCl, 5% Glycerol with EDTA-free protease inhibitors) containing 100 mM, 200 mM, 300 mM, and then 400 mM imidazole. The eluted proteins were checked using SDS-PAGE and pure elutions were combined and passed through Zeba spin desalting 7K MWCO columns (#89892, ThermoFisher Scientific) for desalting. Purified proteins were stored in storage buffer (20mM Tris pH 8.0, 150mM NaCl, 20% Glycerol) at −80 °C. These proteins were sent to Epicypher for further binding analysis.

### Captify™ peptide and nucleosome binding assays

The assay previously known as dCypher™ is now named Captify™, with no distinction in how the assay is performed or its capabilities. Assays were performed as described previously with minor modifications^57–60^. Briefly, 5 µL of Target (biotinylated histone peptides, 100 nM; or biotinylated nucleosomes, 10 nM; *EpiCypher*) was incubated with 5 µL of Query for 30 minutes at 23°C. Interactions were detected by sequential addition of 5 µL Nickel-chelate Acceptor beads (X µg/mL, 30 minutes at 23°C) followed by 5 µL Streptavidin Donor beads (X µg/mL, 30 minutes at 23°C in the dark) prior to measurement of Alpha Counts on a 2104 EnVision Plate Reader (*Revvity*).

This study uses a nomenclature recently devised for accurate scientific communication in the chromatin and epigenetic fields^61^. Here ([H3K4me3]_2_) indicates a fully defined semi-synthetic homotypic nucleosome where other positions not denoted are understood to be definitively unmodified on major histones. For peptide binding curves, Queries were titrated in an optimized peptide binding buffer (50 mM Tris pH 7.5, 25 mM NaCl, 0.01% Tween-20, 0.01% BSA). For nucleosome assays, Queries were first cross-titrated against salt conditions (25 – 200mM NaCl in 25 mM increments) in base nucleosome binding buffer (20 mM Tris, pH7.5, 0.01% NP-40, 1mM DTT, and 0.01% BSA) using ([H3.1]_2_) unmodified, ([H3.1K14ac]_2_), ([H3.1K4acK9acK14acK18ac]_2_), and ([H3.1K4me3K9acK14acK18ac]_2_) nucleosomes to determine optimal probing conditions for a nucleosome discovery screen. Optimal binding conditions were based on the relative EC_50_ (EC_50_^rel^) values for H3K14ac nucleosome binding for each Query: 500 nM BD2 and 25mM NaCl; 107 nM BD3 and125mM NaCl; 316 nM BD4 and 75mM NaCl; 1000 nM BD5 and 25mM NaCl; 33 nM tandem BD2-5 and 150mM NaCl. These optimized binding conditions were then used to determine EC_50_^rel^ values for each BD against a panel of modified nucleosomes selected from discovery screen hits. Data were analyzed using 4-paremeter logistic non-linear regression in GraphPad Prism 10.

### Patient Data

Comparison of PBRM1 mRNA expression between normal breast tissue and tumor tissue was performed in GEPIA using TCGA data. Kaplan Meier plots were generated using Km plotter^62^.

### ChIP-seq data processing of published datasets

Dpf2 and Pbrm1 ChIP-seq datasets from GSE249211^63^ were imported into Partek Flow (v11.0.24.0325). Reads were aligned to mouse genome mm10 build using Bowtie2 (v2.2.5) allowing 1 mismatch. Alignments were filtered for MAPQ>10 using Partek Flow Filter alignments (v11.0.24.0325). Filtered BAMs were used for bigwig generation using deeptools^56^ bamcoverage with --normalizeUsing CPM -p 32 –effectiveGenomeSize 2308125299 -- extendReads. Peak calling was done using MACS (v3.0.0a7) in Partek Flow (v11.0.24.0325) with default parameters and -broad ON mode. Irf1 ChIP-seq datasets from GSE141501^64^ were imported into Partek Flow (v10.0.22.1111), reads were trimmed for quality>20 and minimum length>20 using Trim bases in Partek Flow (v10.0.22.1111). Trimmed reads were aligned to mouse genome mm10 build using Bowtie2 (v2.2.5) allowing 1 mismatch. BAMs were used for bigwig generation using deeptools^56^ bamcoverage with --normalizeUsing CPM -p 32 --effectiveGenomeSize 2494787188. Peak calling was done using MACS (v3.0.0a7) in Partek Flow (v10.0.22.1111) with default parameters. Snail and slug ChIP-seq datasets from GSE61198^65^ were imported into Partek Flow (v12.6.0). Reads were aligned to mouse genome mm10 build using Bowtie (v1.0.0) with default parameters. Alignments were filtered for MAPQ>10 and blacklisted regions using samtools^66^ and bedtools^39^. Filtered BAMs were used for bigwig generation using deeptools^56^ bamcoverage with --normalizeUsing CPM -p 32 --effectiveGenomeSize 2308125299 --extendReads 200 parameters.

## Results

### 1. Pbrm1-depleted cells have reduced capacity to undergo EMT

EMT confers migration and invasion capabilities to epithelial cells^67^ facilitating processes like metastasis. In our previous publication^24^, we found that depletion of *Pbrm1* in epithelial cells resulted in a slight increase in proliferation and a slight reduction in epithelial organization. Further, E-Cadherin expression was slightly decreased, leading us to hypothesize that loss of *Pbrm1* facilitates epithelial-mesenchymal transition (EMT). To test this hypothesis, we treated the normal murine mammary gland (NMuMG) cells with TGFβ1, a cytokine used to induce robust EMT^68^. Five days of TGFβ1 treatment resulted in a morphology change from tightly packed, cobblestone-like epithelial cells to extended, spindle-like mesenchymal cells (**Fig 1A**) which display increased invasion *in vitro* (**Fig 1B,C; SI1A**). While cells transduced with lentiviral-mediated short hairpin RNA (shRNA) against *Pbrm1* initially appeared to lose their epithelial morphology and gain mesenchymal characteristics (**Fig 1A**), contrary to our hypothesis, they did not show accelerated EMT. In fact, *Pbrm1* depleted cells failed to transition to a mesenchymal state upon addition of TGFβ1, as demonstrated by compromised invasive potential (**Fig 1B,C**) and an inability to induce expression of the mesenchymal marker vimentin (**Fig 1D**). After removing TGFβ1 for 2d, E-Cadherin levels in cells with sh*Pbrm1* began to rise again, while E-Cadherin levels stayed repressed in control cells, substantiating their inability to transition to a mesenchymal state (**SI 1B**). This was accompanied by significantly reduced cell viability in sh*Pbrm1* (**Fig 1E**) or sg*Pbrm1* (**Fig 1F**) cells treated with TGFβ1. Similarly, cells with sh*Brd7*, a subunit of the PBAF chromatin remodeling complex required for Pbrm1 incorporation and stability within the complex^4^, displayed a reduction in cell numbers upon TGFβ1 treatment (**SI 1C-E**).

**Figure 1:**
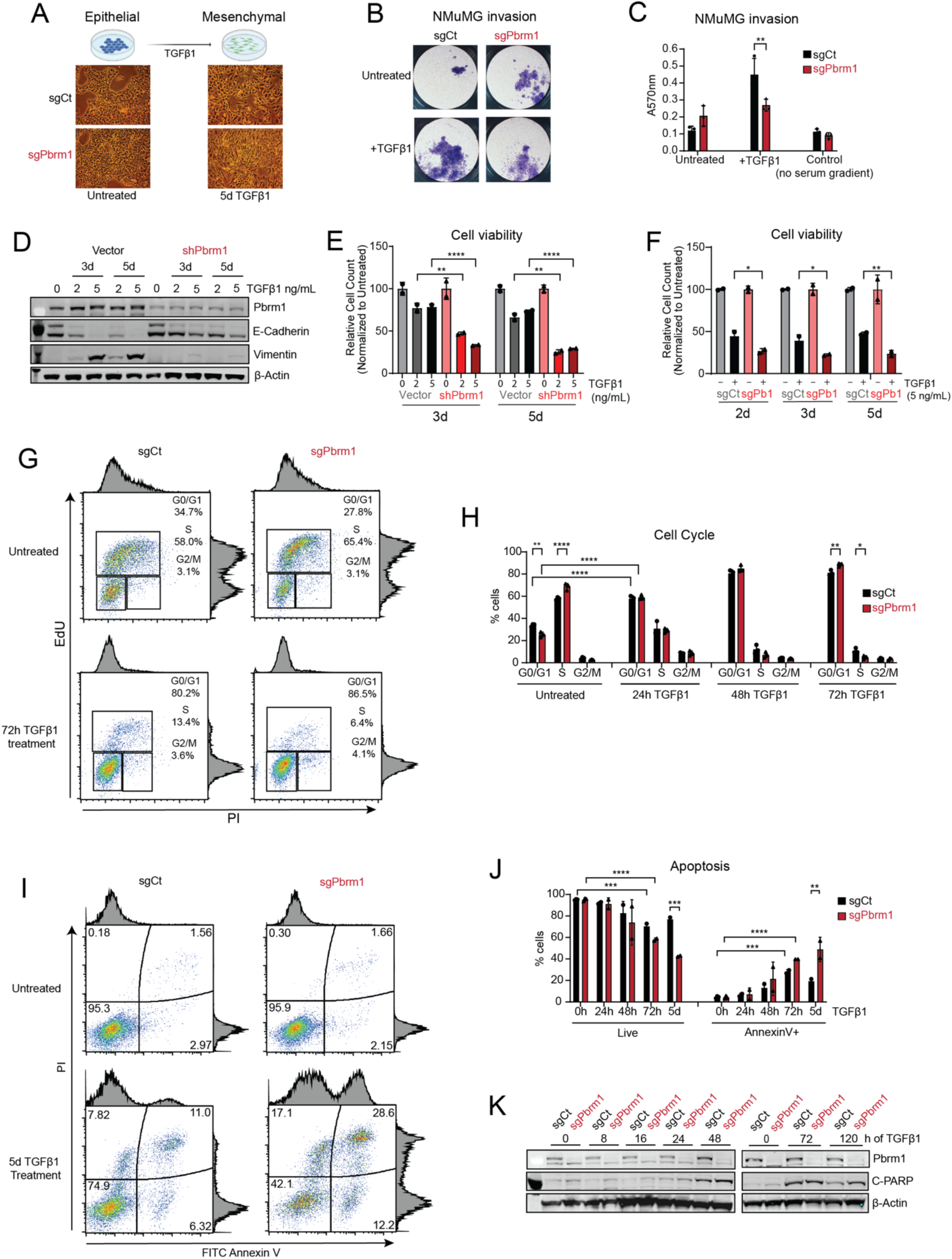
Pbrm1 is important for epithelial-mesenchymal transition in mouse mammary epithelial cells. (A) Schematic of NMuMG cells treatment with TGFβ1 and micrographs of associated morphological changes (4X magnification). (B and C) Representative transwell invasion assay images (B) and bar plot (C) of absorbance quantification of NMuMG sgCt and sg*Pbrm1* cells with and without TGFβ1 treatment. n=3 technical replicates. Data are represented as mean ± SD. (D) Immunoblots of Pbrm1, E-Cadherin, and Vimentin in lysates of NMuMG vector control and sh*Pbrm1* cells with TGFβ1 treatment at the indicated concentrations and incubation times. (E) Relative cell counts of NMuMG control and sh*Pbrm1* cells with and without TGFβ1 treatment, normalized to untreated cells. Representative graph, n=3 biological replicates. Data are represented as mean ± SD. (F) Relative cell counts of NMuMG sgCt and sg*Pbrm1* cells with and without TGFβ1 treatment, normalized to untreated cells. Representative graph, n=3 biological replicates. Data are represented as mean ± SD. (G) Representative flow cytometry density dot plots of Edu and PI staining in fixed NMuMG sgCt and sg*Pbrm1* cells with and without 72h TGFβ1 treatment. The gating strategy for cells in different cell cycle stages (G0/G1, S, or G2/M) is indicated with boxes. (H) Bar plot of the percentage of cells in different cell cycle stages in NMuMG sgCt and sg*Pbrm1* cells with TGFβ1 treatment for the indicated incubation times. Gating was performed as in (G). n=3 biological replicates. Data are represented as mean ± SD. (I) Representative flow cytometry density dot plots of AnnexinV and PI staining in unfixed NMuMG sgCt and sg*Pbrm1* cells with and without 5d TGFβ1 treatment. The gating strategy for the percentage of live (PI-)/dead (PI+) and apoptotic (Annexin V+)/non-apoptotic (Annexin V-) cells is indicated with quadrants. (J) Bar plot of percentage of PI-live cells (left) and Annexin V+ apoptotic cells (right) in NMuMG sgCt and sg*Pbrm1* cells with TGFβ1 treatment for the indicated incubation times. Gating was performed as in (I). n=3 biological replicates. Data are represented as mean ± SD. (K) Immunoblots of Pbrm1 and Cleaved-PARP (C-PARP) from lysates of NMuMG sgCt and sg*Pbrm1* cells with and without TGFβ1 treatment for the indicated time periods. Statistical comparison was done using multiple unpaired t-tests with Holm-Sidak correction. *: p < 0.05, **: p < 0.01, ***: p < 0.001, ****: p < 0.0001

### 2. Pbrm1-depleted epithelial cells exhibit different kinetics of cell cycle and apoptosis under TGFβ1 treatment *in vitro*

To define whether the reduction in cell numbers in *Pbrm1*-depleted cells with TGFβ1 treatment is due to a reduction of proliferation or increased cell death, we evaluated cell cycle and apoptosis. In untreated sg*Pbrm1* cells, there was a higher percentage of cells in S phase, which is consistent with the higher growth rate displayed by sg*Pbrm1* cells (**Fig 1G, H**). In normal epithelial cells, TGFβ1 treatment increases the proportion of cells in G1, which is required to complete EMT^69^. Upon TGFβ1 treatment, sg*Pbrm1* cells initially responded similarly to sgControl (sgCt) cells, with an increase in the percentage of cells in G0/G1 after 24h of treatment; however, with extended TGFβ1 treatment, sg*Pbrm1* cells display a near complete exit from the cell cycle, while sgCt cells continued proliferating (**Fig 1G, H; SI 1F**).

TGFβ1 treatment also induces apoptosis in the subset of NMuMG epithelial cells in G2/M until the cells become mesenchymal^69^. To assess apoptosis in sg*Pbrm1* cells under TGFβ1 treatment, we used cleaved PARP and Annexin V-PI staining. Apoptosis was observed at a similar rate in both groups until 48h of treatment (**Fig 1J**). After 48h, there were significantly higher rates of apoptosis in sg*Pbrm1* cells based on both elevated cleaved-PARP levels and Annexin V staining (**Fig 1I-K; SI 1G**). Thus, with extended TGFβ1 treatment (4-5 days), the sg*Pbrm1* cells exhibited both a cell cycle block and significantly higher cell death relative to control cells.

### 3. Transcriptional profiling of NMuMG cells with TGFβ1 treatment

The induction of EMT is progressive, occurring over several days. Previous work has characterized the early changes in gene expression and chromatin accessibility in NMuMG epithelial cells upon TGFβ1 treatment (2h and 12h)^70^. Since *Pbrm1* depletion affects cells during later stages of EMT (**Fig 1D-K; SI 1B**), we evaluated gene expression after 24, 48 and 72h of TGFβ1 treatment compared to untreated cells. At each time point, TGFβ1 robustly regulated many genes (between 3133 and 4898 differentially expressed genes (DEGs)) with p_adj_<0.05, |log2FC|>1.5) (**SI 2A)** with the majority of DEGs common between the time points (**SI 2B**). Although DEGs were often not different between time groups, the kinetics of gene expression changes were, with some transcripts changing slowly over time, and others changing rapidly, either in a permanent or transient manner (**Fig 2A**). To identify how these gene expression changes correlate with changes in chromatin accessibility, we also performed ATAC-seq. TGFβ1 treatment led to significant changes in chromatin accessibility at all the time points (**Fig 2B, SI 2C**) with ∼15-20% of sites completely gained or lost (**SI 2D**). To determine if changes in chromatin accessibility were associated with gene expression, we compared sites of differential accessibility with the changes in gene expression of the nearest gene (**Fig 2C**). Overall, the changes in accessibility correlate significantly with changes in gene expression (**Fig 2D, SI 2E**). Sites with increased accessibility and expression were enriched for transcription factor (TF) motifs for ATF3, RUNX2, SMAD4, TEAD3, which are all associated with breast cancer EMT^71–74^ (**Fig 2E**), and pathways such as cell migration, extracellular matrix organization, and response to TGFβ1 (**Fig 2F)**. Sites associated with decreased accessibility and expression were enriched for motifs for FRA1, FOXM1, HNF4A, KLF5, which are all involved in various aspects of breast epithelium^75–77^ (**Fig 2G**), and pathways such as cell cycle, DNA replication and epithelium development (**Fig 2H**). Overall, our RNA-seq and ATAC-seq datasets capture the gene expression and chromatin architecture changes induced by TGFβ1 in NMuMG cells.

**Figure 2:**
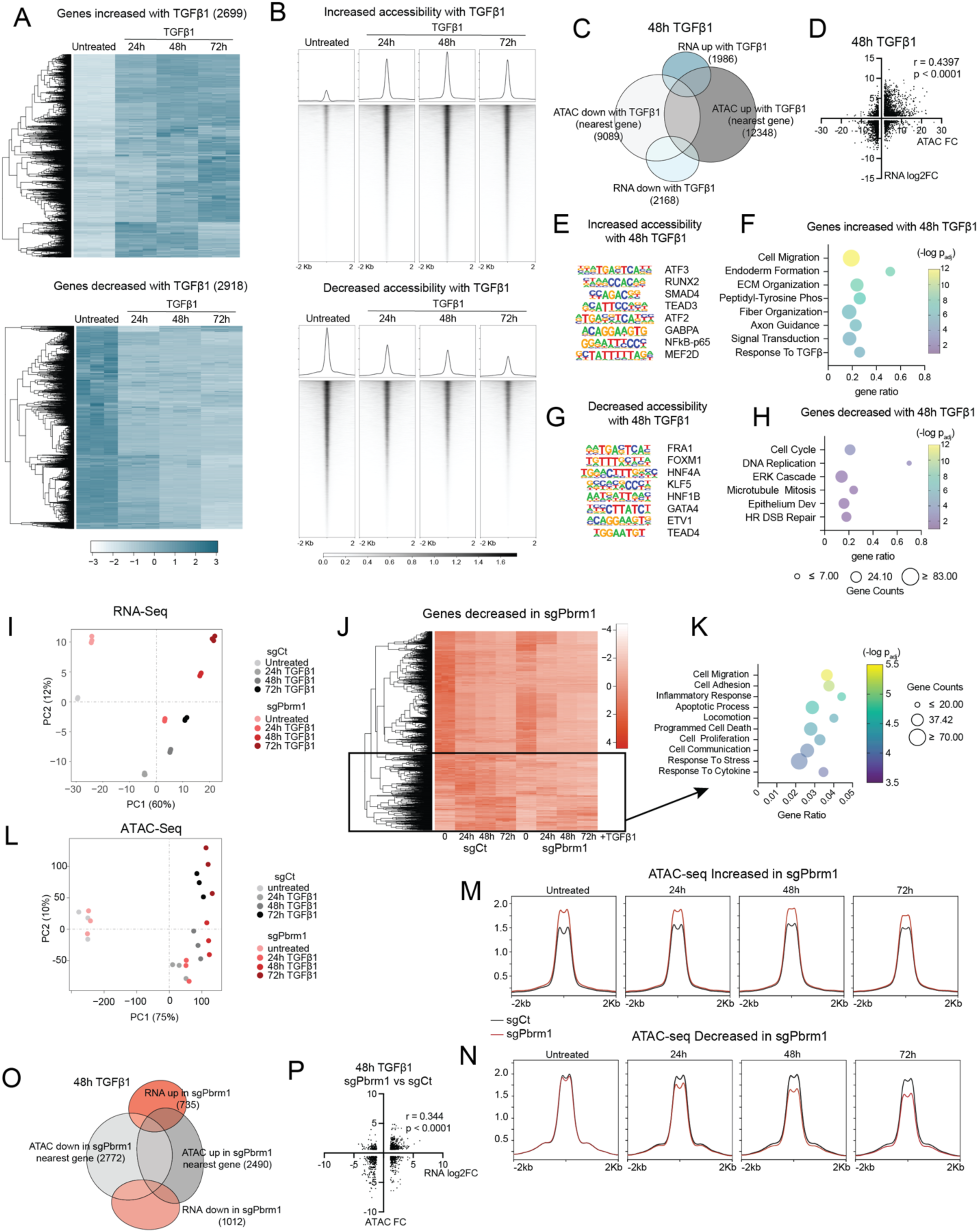
Changes in RNA expression and DNA accessibility upon TGFβ1-mediated EMT in wildtype and Pbrm1 knockout NMuMG cells. (A) Heatmap representation of genes increased (upper) or decreased (lower) in NMuMG sgCt cells with TGFβ1 treatment at different time points. (B) Metagene plots and heatmaps of regions of differential increased (upper) or decreased (lower) accessibility in NMuMG sgCt cells with TGFβ1 treatment. The set of regions includes sites with differential accessibility at any time point of TGFβ1 treatment compared to untreated cells. (C and D) Comparison of changes in RNA expression with changes in accessibility in NMuMG sgCt cells with 48h TGFβ1 treatment relative to no treatment. (C) Venn diagram of overlaps of DEGs from RNA-seq and differentially accessible regions from ATAC-seq annotated to the nearest gene. Total number of genes in each condition is indicated in parentheses in the Venn diagram. (D) Scatter plot of correlation between DEGs from RNA-seq and differentially accessible regions from ATAC-seq annotated to the nearest gene. Each data point in the scatter plot represents the change in expression value of a single gene, *x axis*: differentially accessible regions associated with the nearest gene in ATAC-seq as FC, *y axis*: DEG in RNA-seq as log2FC. The degree of correlation was calculated using all DEGs. (E) Top overrepresented GO terms from pathway analysis using Enrichr on genes with increased expression and accessibility in NMuMG sgCt cells with 48h TGFβ1 treatment. Gene sets were defined using overlap analysis in (C). (F) Motif analysis using Homer on genes with increased expression and accessibility in NMuMG sgCt cells with 48h TGFβ1 treatment. Gene sets were defined using overlap analysis in (C). (G) Top overrepresented GO terms from pathway analysis using Enrichr on genes with decreased expression and accessibility in NMuMG sgCt cells with 48h TGFβ1 treatment. Gene sets were defined using overlap analysis in (C). (H) Motif analysis using Homer on genes with decreased expression and accessibility in NMuMG sgCt cells with 48h TGFβ1 treatment. Gene sets were defined using overlap analysis in (C). (I) Principal Component Analysis of the RNA-seq data of NMuMG sgCt and sg*Pbrm1* cells with and without TGFβ1 treatment for the designated times. (J) Heatmap representation of all genes decreased in NMuMG sg*Pbrm1* relative to sgCt cells at any timepoint of TGFβ1 treatment compared to untreated cells. Genes induced by TGFβ1 in sgCt but not in sg*Pbrm1* cells are subsetted with a box. (K) Top overrepresented GO terms from pathway analysis using Enrichr on the highlighted subset of genes from (J). (L) Principal Component Analysis of the ATAC-seq data for NMuMG sgCt and sg*Pbrm1* cells with and without TGFβ1 treatment where samples are color coded by treatment. (M and N) Metagene plots of all sites of increased (M) or decreased (N) accessibility in NMuMG sg*Pbrm1* cells relative to sgCt cells in any treatment condition. Red lines depict the average enrichment in sg*Pbrm1* cells and black lines depict the average enrichment in sgCt cells. Peak summits are aligned at the center. (O) Venn diagram of DEGs from RNA-seq and the nearest gene of differentially accessible regions from ATAC-seq in sg*Pbrm1* relative to sgCt cells treated with TGFβ1 for 48h. Total number of genes in each condition is indicated in parentheses in the Venn diagram. (P) Scatter plot of differential accessibility from ATAC-seq with the change in expression of the nearest gene in sg*Pbrm1* relative to sgCt cells treated with TGFβ1 for 48h. Each data point in the scatter plot represents the FC expression value of a single gene, *x axis*: DEG in RNA-seq as log2FC, *y axis*: differentially accessible regions associated with the nearest gene in ATAC-seq as FC. Correlation was calculated using all differentially accessible sites.

### 4. Transcriptional profiling of *Pbrm1* knockout cells under TGFβ1 treatment

To understand how Pbrm1 affects TGFβ1 dependent gene expression, we also performed bulk RNA-seq on sg*Pbrm1* NMuMG cells with increasing incubation times with TGFβ1. Principal component analysis (PCA) of the RNA-seq data indicated that cells with sg*Pbrm1* showed distinct gene expression profiles at all time points (**Fig 2I)**. We identified between 1100 and 1800 DEGs (p_adj_<0.05, |log2FC|>1.5) in sg*Pbrm1* cells compared to sgCt, with both shared and unique DEGs across the different treatment conditions (**SI 2F, 2G)**. Most TGFβ1-mediated gene expression changes in sgCt were maintained in sg*Pbrm1* cells; however, there was a subset of ∼300 genes that failed to be induced by TGFβ1 in sg*Pbrm1* cells (**Fig 2J**, highlighted, **SI 2H**). Pathway analysis on these Pbrm1-dependent genes enriched for processes like cell migration, adhesion, inflammatory response, apoptosis, and response to cytokine (**Fig 2K**), consistent with a role for Pbrm1 in facilitating gene expression changes induced by TGFβ1.

### 5. Differential accessibility of *Pbrm1* knockout cells under TGFβ1 treatment

To determine how Pbrm1 contributes to chromatin accessibility during EMT, we also performed ATAC-seq in the sg*Pbrm1* and sgCt cells. Based on PCA, deletion of Pbrm1 had no effect on NMuMG cells under untreated or 24h treatment conditions (**Fig 2L**), and sg*Pbrm1* cells do not even segregate from sgCt cells under these conditions; however, following 48 and 72h TGFβ1 treatment, sg*Pbrm1* and sgCt cells do segregate (**Fig 2L**). To identify changes in accessibility with Pbrm1 loss, we performed MACS analysis and identified 5,000-10,000 sites of differential accessibility (p_adj_<0.05, |FC|>2) in sg*Pbrm1* cells compared to sgCt cells across the different treatments (**SI 2I**). In untreated cells, there were more sites with increased accessibility in sg*Pbrm1* cells than sites with decreased accessibility, as well as slightly higher global accessibility in sg*Pbrm1* cells (**SI 2J**), similar to what has been observed in other cell types^11,27^. In TGFβ1 treated cells, however, there were many more sites with decreased accessibility in the sg*Pbrm1* cells than sites with increased accessibility, particularly at 48 and 72h TGFβ1 treatment (**SI 2I-K**). Metagene analysis of the combined set of sites across treatments with increased or decreased accessibility in sg*Pbrm1* cells compared to sgCt cells, indicates that the magnitude of increased accessibility is consistent across treatments (**Fig 2M**) while the magnitude of decreased accessibility is much higher with TGFβ1 treatment (**Fig 2N**). At 48h TGFβ1 treatment, there is an overlap between gene expression and accessibility changes (**Fig 2O**) as well as a significant positive correlation between DEGs and changes in accessibility at neighboring sites (r =0.344, p<0.0001) (**Fig 2P**). Taken together, these analyses clearly demonstrate that Pbrm1 is necessary for completion of EMT.

### 6. Genome occupancy of PBAF in NMuMG cells under TGFβ1 treatment

To define which of the TGFβ1-regulated genes are direct targets of Pbrm1, we next sought to identify genomic sites bound by PBAF under TGFβ1 treatment conditions. To do this we performed ChIP-seq for Phf10, a subunit of PBAF with high quality ChIP antibodies, along with ChIP-seq for Smarca4, the ATPase subunit present in all BAF subcomplexes (**Fig 3A**). Using MACS, we identified 11,286 robust peaks (**SI 3A**). We compared these sites with published NMuMG ChIP-seq for Pbrm1and a subunit unique to cBAF complexes, Dpf2^63^. As expected, we observed a good overlap between Phf10 peaks and Pbrm1 peaks (∼50% of Pbrm1 sites), moderate overlap with Smarca4 (∼25% of Smarca4 sites), and poor overlap with Dpf2 (∼15% of Dpf2 sites) (**Fig 3B**). In accordance with their presence in different complexes, Phf10 sites displayed low Dpf2 enrichment and vice versa, while Smarca4 was enriched at both subsets of sites (**Fig 3C**). Similar to data from other cell lines^27,78^, Phf10 and Pbrm1 (PBAF) were primarily localized at promoters, while Dpf2 (cBAF) was more enriched at intronic and intergenic sites, and Smarca4 was found at both^79^ (**Fig 3D**). Confirming the enrichment of PBAF at promoters, Phf10 sites showed high enrichment of H3K4me3 and H3K27ac, and low enrichment of H3K4me1 histone modifications (**Fig 3E**).

**Figure 3:**
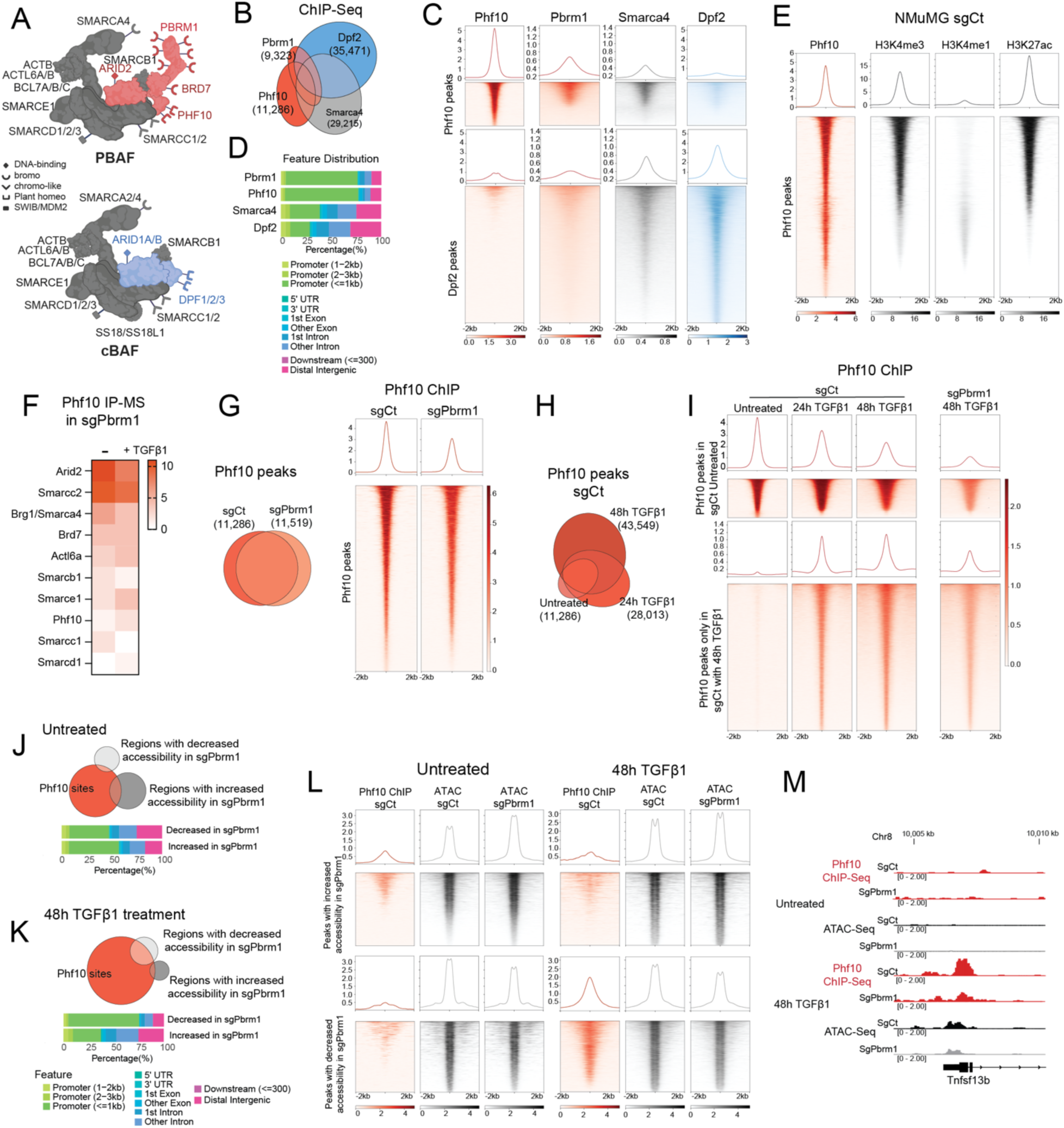
PBAF is recruited to new sites upon TGFβ1 treatment in NMuMG cells to increase DNA accessibility. (A) Schematic representation of the PBAF and cBAF complexes with shared subunits in grey, PBAF-specific subunits in red, and cBAF specific subunits in blue. The reader domains encoded by the different subunits and their respective functions are indicated. (B) Venn diagram of Phf10 and Smarca4 ChIP-seq peaks from NMuMG sgCt cells with Pbrm1 and Dpf2 ChIP-seq peaks from NMuMG cells (GSE249211). Total number of peaks identified in ChIP-seq for each protein is indicated in parentheses. (C) Metagene plots and heatmaps of ChIP-seq enrichment of Phf10, Pbrm1, Smarca4, and Dpf2 at Phf10 and Dpf2 ChIP-seq sites in NMuMG cells as described in (B). (D) Genomic feature distribution of the ChIP-seq peaks identified for Pbrm1, Phf10, Smarca4, and Dpf2 in NMuMG cells as described in (B). (E) Metagene plots and heatmaps of ChIP-seq enrichment of Phf10 H3K4me3, H3K4me1, and H3K27ac enrichment at Phf10 ChIP-Seq sites in sgCt cells as described in (B). (F) Heatmap representation of PBAF subunit abundance in NMuMG sg*Pbrm1* cells with and without TGFβ1 treatment identified by Phf10 IP-MS. (G) (left)Venn diagram of Phf10 ChIP-seq peaks in NMuMG sgCt and sg*Pbrm1* cells. Total number of Phf10 ChIP-seq peaks identified in each condition is indicated in parentheses. (right) Metagene plots and heatmaps of ChIP-seq enrichment of Phf10 in sgCt and sg*Pbrm1* cells at Phf10 ChIP-seq sites in sgCt cells as described in (B). Peak summits are aligned at the center. (H) Venn diagram of Phf10 ChIP-seq peaks in untreated and TGFβ1-treated NMuMG sgCt cells. Total number of Phf10 ChIP-seq peaks identified in each condition is indicated in parentheses. (I) Metagene plots and heatmaps of Phf10 ChIP-seq enrichment in untreated and TGFβ1-treated NMuMG cells. Enrichment was plotted at Phf10 sites in untreated sgCt cells (top) and Phf10 sites unique to TGFβ-treated sgCt cells. Peak summits are aligned at the center. (J) (top) Venn diagram of Phf10 ChIP-seq peaks in NMuMG sgCt cells and sites of differential accessibility in sg*Pbrm1* cells compared to sgCt cells. (bottom) The genomic feature distribution for sites of differential accessibility in sg*Pbrm1* cells compared to sgCt cells (K) (top) Venn diagram of Phf10 ChIP-seq peaks in NMuMG sgCt cells with 48h TGFβ1 treatment and sites of differential accessibility in sg*Pbrm1* cells compared to sgCt cells with 48h TGFβ1 treatment. (bottom) The genomic feature distribution for sites of differential accessibility in sg*Pbrm1* cells compared to sgCt cells with 48h TGFβ1 treatment. (L) Metagene plots and heatmaps of Phf10 ChIP-seq and ATAC-seq enrichment at sites of differential accessibility in sg*Pbrm1* cells compared to sgCt cells under untreated (left) and 48h TGFβ1-treated (right) conditions. (M) Genomic tracks of Phf10 ChIP-seq and ATAC-seq enrichment at the *Tnfsf13b* locus in sgCt and sg*Pbrm1* cells in untreated (top) and 48h TGFβ1-treated (bottom) conditions.

Pbrm1 is the last subunit to be incorporated into PBAF^4^ and as such, PBAF complexes can form and exist in the absence of Pbrm1^80^. Confirming this, IP-MS of PBAF subunit Phf10 in sg*Pbrm1* cells identified all PBAF subunits in both untreated and TGFβ1 treated cells (**Fig 3F**). Therefore, we used Phf10 ChIP-seq to determine how loss of Pbrm1 affects PBAF chromatin binding. There was a high overlap between Phf10 peaks from sgCt and sg*Pbrm1* cells (**Fig 3G**), indicating that PBAF retains binding to chromatin in the absence of Pbrm1; however, there was a decrease in Phf10 enrichment in sg*Pbrm1* cells, which was confirmed by qPCR in an independent replicate (**Fig 3G; SI3B**). This is in agreement with our previous findings that Pbrm1 loss decreases, but doesn’t abrogate, PBAF affinity to chromatin^78,80^.

To identify how PBAF localization changes during EMT, we next performed Phf10 ChIP-seq at 24 and 48h TGFβ1 treatment. The majority of the Phf10 sites identified in untreated NMuMG cells were maintained across conditions; however, a large number of additional peaks appeared with 24 and 48h of TGFβ1 treatment (**Fig 3H**, **SI 3C),** which were globally reduced in sg*Pbrm1* cells (**Fig 3I; SI 3D, E**), These findings indicate that Pbrm1 plays a critical in PBAF redistribution during EMT to TGFβ1-inducible sites.

### 7. Integration of accessibility changes in Pbrm1-depleted cells with Phf10 binding

Next, we evaluated how PBAF binding relates to changes in accessibility in sg*Pbrm1* cells. In untreated cells, Phf10 is not highly enriched at sites with changes in accessibility with sg*Pbrm1* (**Fig 3J**). With 48h TGFβ1 treatment, however, Phf10 peaks sites do overlap with (**Fig 3K**) and are enriched at (**Fig 3L**) sites of decreased accessibility in sg*Pbrm1* cells. Collectively, these data suggest that TGFβ1-inducible promoters such as *Tnfsf13b*, a cytokine associated immune escape and poor prognosis in cancer^81,82^ requires Pbrm1/PBAF for maximal accessibility (**Fig 3M**) while constitutive, active promoters do not require Pbrm1 for significant changes in accessibility (**SI 3F**, **G**)

### 8. PBAF cooperates with inducible transcription factors during EMT

To identify the TFs that cooperate with PBAF for transcription, we performed DiffTF^52^ analysis on the regions of accessibility identified in our ATAC-seq. Regions that required Pbrm1 for increased accessibility upon TGFβ1 treatment were enriched for AP-1 motifs (Jun, Junb, Jund, Fosb, Fosl1, Fosl2), EMT TFs (Snai1, Snai2, Zeb1), SWI/SNF (Smarcc1) and methyl CpG binding protein 2 (**Fig 4A**). cBAF has been demonstrated to cooperate with the AP-1 family of TFs at enhancers^83,84^; however, this has not been demonstrated for PBAF at promoters. To elucidate the AP-1 subunits specifically important for TGFβ1-inducible gene expression, we used our RNA-seq datasets to identify which AP-1 subunit genes are induced by TGFβ1. The expression of *Jun*, *Junb*, *Fosl2*, *Fosb* and *Atf3* were all significantly increased upon TGFβ1 treatment (**SI 4A**). We next performed ChIP-seq for these five TFs and obtained peaks for Atf3, Fosb and Fosl2. For all three of these AP-1 subunits, enrichment was much higher after 48h of TGFβ1 treatment, confirming their inducible expression and function during EMT (**Fig 4B**). Atf3 and Fosb had reduced enrichment in sg*Pbrm1* cells; in contrast, Fosl2 enrichment was increased (**Fig 4B, SI 4B**) consistent with data from melanoma cells with loss of PBAF^27^. To identify whether any of the AP-1 subunits directly cooperate with PBAF, we performed overlap analysis on the genomic regions identified for each protein. Only Atf3 peaks had high overlap and co-enrichment with Phf10 peaks (**Fig 4C**). Supporting this co-enrichment, Atf3 peaks were preferentially enriched at promoters, like Phf10 peaks, while Fosb and Fosl2 peaks were more enriched at intronic and intergenic regions (**Fig 4D**). Many of the inducible Phf10 sites, such as the promoter of *Tnfsf13b* (**Fig 3M**), have similar profiles for Phf10 and Atf3 binding (**Fig 4E**), supporting Atf3’s dependency on PBAF for chromatin binding.

**Figure 4:**
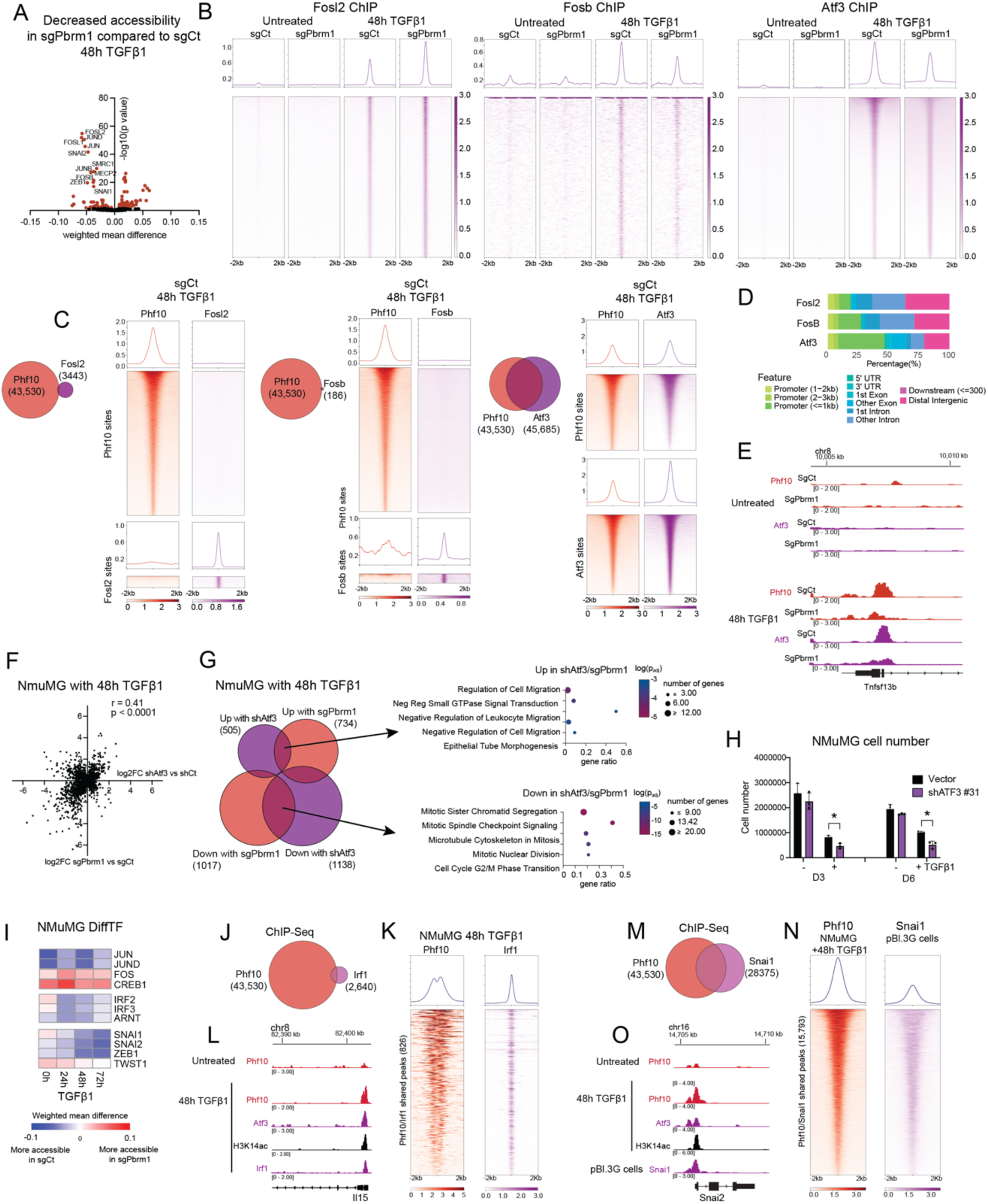
Pbrm1 facilitates inducible transcription factors upon TGFβ1 treatment. (A) Volcano plot of TF consensus motifs enriched in regions of differential accessibility in NMuMG sg*Pbrm1* relative to sgCt cells with 48h TGFβ1 treatment (FDR<0.001). (B) Metagene plots and heatmaps of ChIP-seq enrichment of Fosl2 (left), Fosb (center) and Atf3 (right) in sgCt and sg*Pbrm1* cells in both untreated and 48h TGFβ1-treated conditions. Enrichment was plotted for corresponding TF binding sites compiled from all conditions. (C) Venn diagram of Phf10 ChIP-Seq sites with Fosl2 (left), Fosb (center) and Atf3 (right) ChIP-seq sites in sgCt cells with 48h TGFβ1 treatment. Total number of peaks identified in ChIP-seq for each protein is indicated in parentheses. To the right of each Venn diagram are the metagene plots and heatmaps of ChIP-seq enrichment of Phf10 with Fosl2 (left), Fosb (center) or Atf3 (right) at the Phf10 (top) and associated TF (bottom) binding sites. (D) Genomic feature distribution of the Fosl2, Fosb, and Atf3 ChIP-seq peaks. (E) Genomic tracks of Phf10 and Atf3 ChIP-seq enrichment in sgCt and sg*Pbrm1* cells at *Tnfsf13b* locus in untreated (top) and 48h TGFβ1-treated (bottom) conditions. (F) Scatter plot of the change in expression for DEGs from sg*Pbrm1* vs sgCt with 48h TGFβ1 plotted against DEGs from sh*Atf3* vs shScr with 48h TGFβ1. Each data point in the scatter plot represents the log2FC expression value of a single gene in the indicated comparison, *x axis*: sh*Atf3* vs shCt, *y axis*: sg*Pbrm1* vs sgCt. The degree of correlation was calculated using all DEGs. (G) Venn diagram of overlap between DEGs in sg*Pbrm1* vs sgCt and DEGs from sh*Atf3* vs shScr, both with 48h TGFβ1 treatment. Total number of genes from each condition is indicated in parentheses. The top overrepresented GO terms for the set of genes increased by both sg*Pbrm1* and sh*Atf3* (top) or decreased by both sg*Pbrm1* and sh*Atf3* (bottom) were identified using Enrichr pathway analysis. (H) Absolute cell counts of control and sh*Atf3* cells with and without 5 ng/mL TGFβ1 treatment. 0.9 million cells were seeded for each cell line and cell counts were taken on day 3 and 6. Representative graph, n=3 biological replicates. Data are represented as mean ± SD. (I) Heatmap representation of the differential accessibility of consensus TF motifs in NMuMG sg*Pbrm1* cells with and without TGFβ1 treatment performed using diffTF analysis. Significant differences in weighted means between the groups are shown. (J) Intersection of Phf10 ChIP-seq peaks in NMuMG sgCt cells with Irf1 ChIP-seq peaks in 48h TGFβ1-treated NMuMG cells from a published Irf1 ChIP-Seq dataset (GSE141501). Total number of peaks identified in ChIP-seq for each protein is indicated in parentheses. (K) Metagene plots and heatmaps of ChIP-seq enrichment of Phf10 from 48h TGFβ1-treated NMuMG sgCt cells and Irf1 from 48h TGFβ1-treated NMuMG cells from a published dataset at Phf10 and Irf1 shared sites. (L) Genomic tracks of ChIP-seq enrichment of Phf10, Atf3, H3K14ac, and Irf1 in untreated and 48h-TGFβ1 treated NMuMG sgCt at *Il15* locus. (M) Intersection of Phf10 ChIP-seq peaks in NMuMG sgCt cells with Snai1 ChIP-seq peaks in pBI3G mouse mesenchymal breast cancer cells from a published dataset (GSE61198). Total number of peaks identified in ChIP-seq for each protein is indicated in parentheses. (N) Metagene plots and heatmaps of ChIP-seq enrichment of Phf10 from 48h TGFβ1-treated NMuMG sgCt cells and Snai1 from pBI3G mouse mesenchymal breast cancer cells at Phf10 and Snai1 shared sites. (O) Genomic tracks of ChIP-seq enrichment at *Snai2* locus of Phf10, Atf3, and H3K14ac in untreated and 48h-TGFβ1 treated NMuMG sgCt cells and Snai1 ChIP-Seq enrichment in pBI3G mouse mesenchymal breast cancer cells. *: p < 0.05, **: p < 0.01, ***: p < 0.001, ****: p < 0.0001

To further understand the functional relationship between Atf3 and PBAF, we performed RNA-seq on NMuMG cells depleted for *Atf3* under normal conditions and following 48h TGFβ1 treatment (**SI 4C, D, E**). There was a significant positive correlation between the DEGs in sh*Atf3* and sg*Pbrm1* cells treated with TGFβ1 (r=0.41, p<0.0001) (**Fig 4F**), with many overlapping DEGs (**Fig 4G**). Pathway analysis on genes increased by sh*Atf3* and sg*Pbrm1* under TGFβ1-treatment enriched for negative regulation of cell migration and epithelial tube morphogenesis, while the genes decreased by sh*Atf3* and sg*Pbrm1* enriched for processes related to cell cycle. To determine if Atf3 is also required in NMuMG cells during EMT, we assessed cell viability upon TGFβ1 treatment. sh*Atf3* cells exhibited lower cell survival relative to shCt, like cells with *Pbrm1* depletion (**Fig 4H**). These results were consistent between two separate hairpins against Atf3 (**SI 4F**). Overall, these results reflect a role for Atf3 in proliferation and viability during EMT.

The data above strongly suggest that Atf3 is dependent on Pbrm1 for chromatin binding (**Fig 4B-E, SI 4B**). However, Atf3 is a stress-inducible TF upregulated by TGFβ1 treatment^71,85,86^ and its induction was reduced by *Pbrm1* deletion (**SI 4A**). Like *Atf3* mRNA expression levels, TGFβ1-induced protein levels of Atf3 were also blunted upon deletion of *Pbrm1* (**SI 4G**). Therefore, even though Atf3 co-binds with PBAF across the genome, the reduction in Atf3 binding upon *Pbrm1* deletion could be due to decreased Atf3 protein levels. To evaluate whether the decrease in Atf3 expression in sg*Pbrm1* cells is solely responsible for the observed phenotypes, we overexpressed human ATF3 in sgCt and sg*Pbrm1* cells (**SI 4H**). Atf3 overexpression did not rescue the cell viability defect in sg*Pbrm1* cells upon TGFβ1 treatment, suggesting that PBAF plays a role in both Atf3 regulation as well as function in NMuMG cells (**SI 4I**). Atf3, like other AP-1 subunits, requires cooperation with additional TFs to regulate gene expression^83^. To identify potential cell-type specific TFs that cooperate with Atf3 and PBAF to regulate TGFβ1-dependent genes in NMuMG cells, we again turned to the DiffTF data to identify consensus TF sites with reduced accessibility specifically in TGFβ1-treated sg*Pbrm1* cells. Unlike AP-1 sites that have decreased (Jun) or increased (Fos) accessibility in both untreated and treated sg*Pbrm1* cells, consensus TF sites associated with Irf and Snail TFs were decreased in sg*Pbrm1* cells only when treated with TGFβ1 (**Fig 4I**). The IRF family of TFs are established mediators of IFN signaling pathways^87^, and the Snail family primarily mediates signals originating from TGFβ1 stimulation^88^. To identify which specific members of these two families are involved in EMT in NMuMG cells, we again turned to the RNA-seq datasets. Of the nine Irf family members, only Irf1 is induced by TGFβ1 in NMuMG cells (**SI 4J**). Irf1 was previously reported to be necessary for EMT in NMuMG cells^64^. Knockdown of *Irf1* increased proliferation in untreated NMuMG cells but decreased viability under TGFβ1 treatment^64^, which is consistent with our observations in sh*Pbrm1* cells. Using published ChIP-seq data of Irf1 in NMuMG cells with 48h TGFβ1 treatment^64^, we identified 2640 Irf1 peaks, of which ∼30% were co-bound by Phf10 (**Fig 4J, K**) including Pbrm1-dependent inducible genes such as *Il15* (**Fig 4L**), *Cxcl10, Jak2,* and *Csf1*.

We similarly used the RNA-seq datasets to define the expression levels of the three Snail TF family members in NMuMG cells during EMT. Only *Snai1* was induced with TGFβ1 in NMuMG cells (**SI 4K**). Using published ChIP-seq data of *Snai1* from mouse mesenchymal pBI.3G breast cancer cells^65^, we identified 28,375 *Snai1* peaks, of which ∼55% were co-bound by Phf10 in TGFβ1-treated NMuMG cells (**Fig 4M, N**), including many sites with inducible Phf10 recruitment. One of these sites is Snai2, which is repressed by Snai1 at later stages of EMT^89^ (**Fig 4O**). In agreement with the strong overlap between these TFs and Phf10, both Irf1 and Snai1 peaks are also enriched for promoters (**SI 4L**). Collectively, these data provide evidence that PBAF facilitates the binding of TGFβ1-inducible TFs involved in proliferation, survival, migration and immunosuppression during EMT.

### 9. Pbrm1 bromodomains recognize H3K14ac

We next sought to define how PBAF is targeted to specific sites during EMT. We previously demonstrated that only singly modified H3_1-30_K14ac and K14ac-containing multiply acetylated peptides (H3_1-30_K14acK18acK23acK27ac) were capable of enriching the PBAF complex from nuclear lysates^80^. To evaluate Pbrm1’s contribution, we expressed recombinant BDs 2, 3, 4, and 5, and tandem BDs 2-5 for *in vitro* analysis (**Fig 5A**). Previous studies indicate BD2-5 are the critical bromodomains for Pbrm1 interactions in cells^80^ and *in vitro*^90^. Using Captify™ (**Fig 5B)**, peptide binding assays revealed that none of the BDs bound tightly to H3_1-20_K14ac alone; however, all queries except BD5 bound to peptides containing multiple acetylations (**Fig 5C, SI 5A**). with the highest affinity interaction between tandem BD2-5 and H3_1-20_K4acK9acK14acK18ac peptides (EC_50_^rel^: ∼8 nM) (**Fig 5D).** Although peptides indicate PBAF/Pbrm1 may recognize multiple acetylations on H3; histone PTM recognition can differ substantially in the nucleosome context^57^, particularly given our recent finding that several Pbrm1 BDs bind nucleic acids^78^. We therefore used Captify™ to screen a collection of designer nucleosomes using individually optimized conditions (see methods). BD2 showed specificity for nucleosomes with K14ac, while the other BDs recognized multiple acetylation marks^91^ (**SI 5B**). EC_50_^rel^ values were calculated for 11 acetylated nucleosomes and an unmodified control (**Fig 5E, SI 5C)**. Not surprisingly, the lowest EC_50_^rel^ values (∼20 nM) were observed for the tandem BD2-5 (**Fig 5E**). Consistent with peptide pulldowns, only nucleosomes with H3K14ac were enriched. Unlike peptide experiments, however, there was very little increase in binding affinity for nucleosomes with multiple acetylation marks compared to H3K14ac alone (**Fig 5F**). This indicates that, in the context of the nucleosome, Pbrm1 primarily recognizes H3K14ac via BD2 with the other bromodomains contributing to overall binding affinity through additional interactions with H3K14ac, nucleic acids, and minimally with other acetylation marks.

**Figure 5:**
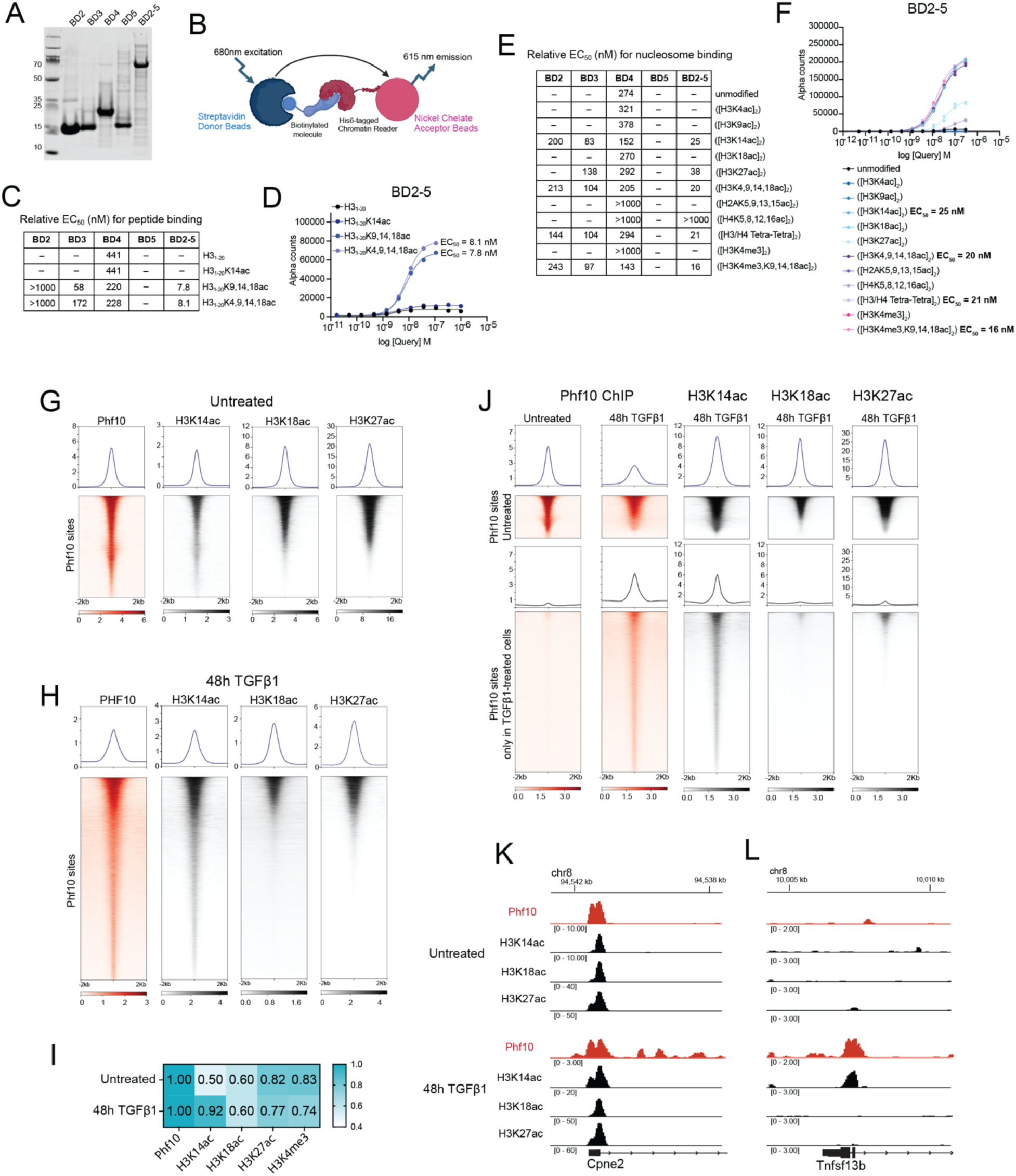
Pbrm1 binds H3K14ac at TGFβ1-inducible PBAF binding sites. (A) Coomassie gel of the purified recombinant proteins BD2, BD3, BD4, BD5 and the tandem BD2-5 used for peptide and nucleosome binding assays. (B) Schematic representation of EpiCypher’s Captify™ assay. (C) Table of EC_50_ values (nM) of the different BDs for the indicated peptides obtained using the ALPHA/dCypher assay. HP1 binding to H3K9me3 peptide is used as a positive control in this assay. (D) Binding curves of tandem BD2-5 with the indicated peptides obtained using the ALPHA/dCypher assay. EC_50_ (nM) values are indicated next to the corresponding curves. (E) Table of the relative EC_50_ values (nM) of the different BDs for nucleosomes bearing the indicated histone modifications obtained using the ALPHA/dCypher assay. HP1 binding to H3K9me3 nucleosomes is used as a positive control in this assay. (F) Binding curves of tandem BD2-5 for nucleosomes bearing the indicated peptides obtained using the ALPHA/dCypher assay. HP1 binding to H3K9me3 peptide is used as a positive control in this assay. EC_50_ values (nM) obtained for positive binders are indicated in the legend. (G and H) Metagene plots and heatmaps of ChIP-seq enrichment of Phf10, H3K14ac, H3K18ac, and H3K27ac at Phf10 binding sites in untreated (H) and 48h TGFβ1-treated (I) sgCt cells. (I) Correlation matrix with r-values between Phf10 and H3K14ac, H3K18ac, H3K27ac, and H3K4me3 ChIP-seq enrichment in untreated and 48h TGFβ1-treated sgCt cells. (J) Metagene plots and heatmaps of ChIP-seq enrichment of Phf10, H3K14ac, H3K18ac, and H3K27ac in untreated and 48h TGFβ1-treated sgCt cells. The top heatmap is at Phf10 binding sites in untreated cells and the bottom is Phf10 binding sites only found in TGFβ1-treated cells. (K and L) Genomic tracks of ChIP-seq enrichment of Phf10, H3K14ac, H3K18ac, and H3K27ac in untreated (M) and 48h-TGFβ1 treated (M) sgCt cells at constitutive locus *Cpne2* and an inducible locus *Tnfsf13b*.

### 10. Pbrm1 facilitates PBAF recruitment to sites of TGFβ1-induced H3K14ac

To elucidate the histone marks recognized by PBAF in a cellular context, we performed ChIP-seq in NMuMG cells for H3K14ac, H3K18ac, and H3K27ac. Consistent with PBAF binding at active promoters, all three histone marks were similarly enriched at Phf10 binding sites (**Fig 5G**); however, H3K14ac was most highly enriched at Phf10 binding sites upon TGFβ1 treatment (**Fig 5H**). This was confirmed by calculating correlation scores between ChIP-seq datasets for histone marks and Phf10. While all three histone acetylation marks, as well as H3K4me3, correlate with Phf10 binding in untreated cells to a similar degree, H3K14ac is more highly correlated that other acetylation marks to Phf10 in TGFβ1 treated cells (**Fig 5I**). We next evaluated which histone acetylation marks are associated with constitutive and TGFβ1-induced Phf10 binding sites. Under TGFβ1 treatment, all three acetylation marks are maintained at constitutive Phf10 sites, while only H3K14ac is at inducible Phf10 sites (**Fig 5J-L**). These findings strongly suggest that Pbrm1 is required for recruiting PBAF to the H3K14ac histone marks at TGFβ1-inducible sites, where it is present in the absence of other H3 acetylation marks.

### 11. PBRM1 in human breast cancer metastasis

Given our findings about Pbrm1 during EMT, we sought to determine whether Pbrm1 is required for metastasis. Pbrm1 was previously implicated as a tumor suppressor in breast cancer^92^, with a low rate of mutation in patients (1.2%)^22^ and a reported decrease in protein levels in cancer tissue compared to normal breast tissue in a small cohort of 150 patients^93^. Using the larger TCGA breast cancer dataset, we found that *PBRM1* expression is slightly higher in breast cancer tissues compared to the normal counterparts (**Fig 6A**). *PBRM1* expression levels were not associated with disease progression in all breast cancers or early-stage breast cancer (**Fig 6B** left, center). In contrast, in metastatic triple negative breast cancer (mTNBC), *PBRM1* expression is highly predictive of worse overall survival (**Fig 6B** right). These findings suggest a context-dependent requirement for *PBRM1* in the metastatic setting. Along these lines, we found that *PBRM1* depletion does not affect the viability of the non-metastatic MCF10CA1h (CA1h) human breast cancer cell line but is required for viability in the isogenic and metastatic MCF10CA1a (CA1a) cell line (**SI6A-C**). cBAF-specific subunit *ARID1A* also showed elevated expression in breast cancer tissues relative to normal group (**Fig 6C**); however, in contrast to *PBRM1* expression, higher *ARID1A* expression was associated with better survival in mTNBC (**Fig 6D**). Based on these data from TCGA patient samples and human cell lines, we hypothesized that *PBRM1* expression may contribute to progression of metastatic breast cancer.

**Figure 6:**
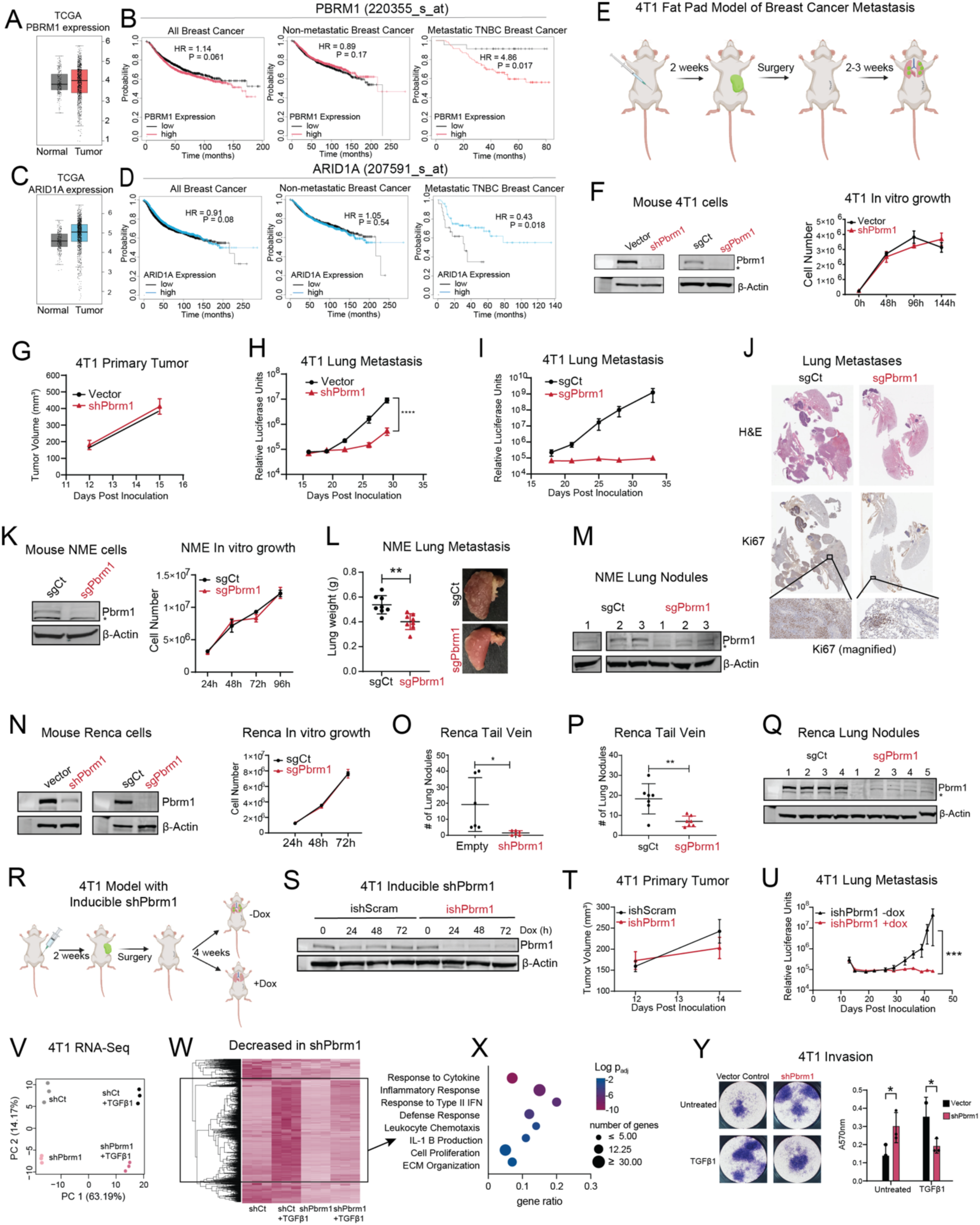
Pbrm1 is required for breast cancer metastasis in mice. (A) Box plot of the mRNA expression of *PBRM1* in normal vs tumor tissues of the TCGA human breast cancer dataset. Generated using GEPIA. (B) Kaplan-Meier curves displaying the effect of *PBRM1* mRNA expression on overall survival in all breast cancer (left), non-metastatic breast cancer (center) and metastatic TNBC (right) patient samples. Curves generated using Km plotter. (C) Box plots of the mRNA expression of *ARID1A* in normal vs tumor tissues of the TCGA human breast cancer dataset. Generated using GEPIA. (D) Kaplan-Meier curves displaying the effect of *ARID1A* mRNA expression on overall survival in all breast cancer (left), non-metastatic breast cancer (center) and metastatic TNBC (right) patient samples. Curves generated using Km plotter. (E) Schematic representation of the syngeneic 4T1 fat pad model of breast cancer metastasis. 4T1 cells expressing luciferase are injected orthotopically into the inguinal fat pad, the primary tumor is surgically removed, and lung metastases are monitored using in vivo imaging. (F) (left) Immunoblots of whole cell extracts of 4T1 cells treated with shRNA or sgRNA against *Pbrm1.* (right) *In vitro* proliferation of 4T1 sh*Pbrm1* and control cells. n=2 biological replicates. Two-way ANOVA was used for statistical comparison. Data are represented as mean ± SD. (G) Primary tumor growth in 4T1 sh*Pbrm1* and control cells measured using vernier calipers at the indicated time points. Two-way ANOVA with multiple comparisons was used for statistical comparison. Data are represented as mean ± SEM. (H-I) Bioluminescent imaging of lung metastasis in 4T1 sh*Pbrm1* and control cells (H) and sg*Pbrm1* and sgCt cells (I) after removal of the primary tumor. Two-way ANOVA with multiple comparisons was used for statistical comparison. Data are represented as mean ± SEM. (J) H&E and Ki-67 stained IHC images of the lungs harvested at the end of the experiment described in (I). (K) (left) Immunoblots of whole cell extracts of NME cells treated with sgRNA against *Pbrm1.* (right) *In vitro* proliferation of NME sg*Pbrm1* and control cells. n=2 biological replicates. Two-way ANOVA was used for statistical comparison. Data are represented as mean ± SD. (L) Scatter plot of lung weights from individual mice in the NME sgCt and sg*Pbrm1* groups harvested at the end of experiment. Welch’s t-test was used for statistical comparison. Data are represented as mean ± SD. Images of representative lung halves bearing metastatic lung nodules from the two groups are shown. (M) Immunoblots of the other lung halves from (L), which were cultured *in vitro* under antibiotic selection. Whole cell extracts were used for immunoblotting. (N) (left) Immunoblots of whole cell extracts of Renca cells treated with shRNA or sgRNA against *Pbrm1.* (right) *In vitro* proliferation of Renca sg*Pbrm1* and control cells. n=3 biological replicates. Two-way ANOVA was used for statistical comparison. Data are represented as mean ± SD. (O and P) Scatter plot of the number of lung nodules from individual mice in the Renca sh*Pbrm1* and control groups (O), and sg*Pbrm1* and sgCt groups (P) harvested at the end of experiment. Welch’s t-test was used for statistical comparison. Data are represented as mean ± SD. (Q) Immunoblots of the lung halves from (P), which were cultured *in vitro* under antibiotic selection. Whole cell extracts were used for immunoblotting. (R) Schematic representation of the syngeneic 4T1 fat pad model from of breast cancer metastasis from (E) using doxycycline-inducible *Pbrm1* KD after primary tumor removal. (S) Immunoblots of whole cell extracts of 4T1 cells after doxycycline-induced expression of shRNA against *Pbrm1.* 2ug/ml doxycycline concentration was used for induction of hairpin expression. (T) Primary tumor growth in 4T1 ish*Pbrm1* and control cells measured using vernier calipers at the indicated time points. Two-way ANOVA with multiple comparisons was used for statistical comparison. Data are represented as mean ± SEM. (U) Bioluminescent imaging of lung metastasis in 4T1 i*shPbrm1* cells with and without doxycycline administration after removal of the primary tumor. Two-way ANOVA with multiple comparisons was used for statistical comparison. Data are represented as mean ± SEM. (V) Principal Component Analysis of the RNA-seq profile of 4T1 shCt and sh*Pbrm1* cells with and without TGFβ1 treatment where samples are color coded by treatment. (W) Heatmap representation of genes increased decreased in 4T1 sh*Pbrm1* cells with and without TGFβ1 treatment. (X) Top overrepresented GO terms from pathway analysis using Enrichr on the highlighted subset of genes from (C). (Y) Transwell invasion assay images and bar plot of absorbance quantification of 4T1 control and sh*Pbrm1* cells with and without TGFβ1 treatment. n=3 technical replicates. Data are represented as mean ± SD. *: p < 0.05, **: p < 0.01, ***: p < 0.001, ****: p < 0.0001

### 12. Pbrm1-depleted cells exhibit reduced metastasis in mouse models

To evaluate the role of Pbrm1 in metastasis *in vivo*, we used the 4T1 model of breast cancer metastasis. 4T1 cells expressing luciferase were orthotopically implanted into the mammary fat pads of syngeneic Balb/c mice. When the primary tumor reached a group average of ∼400 mm^3^ in size (∼2 weeks), it was surgically removed, and the outgrowth of lung metastases was monitored using *in vivo* bioluminescence imaging (**Fig 6E**). *Pbrm1* depletion or deletion did not significantly affect 4T1 cell proliferation *in vitro* (**Fig 6F**) or in the orthotopic primary tumor (**Fig 6G; SI 6D**). In contrast, *Pbrm1* depletion or deletion resulted in a significant reduction of pulmonary metastasis as measured by bioluminescence (**Fig 6H, I**) and end-point necropsy (**Fig 6J**). To determine if this dependency is unique to PBAF, we generated 4T1 cell lines with KD of a second PBAF subunit, *Brd7*, or cBAF-specific subunit, *Arid1A* (**SI 6E**) and similarly evaluated primary tumor growth and metastasis *in vivo*. *Brd7* depletion resulted in a reduction in Pbrm1 protein levels in these cells, as noted earlier (**SI 1B**). There were no significant differences observed in the primary tumor growth with sh*Brd7* or sh*Arid1a* (**SI 6F, G**). In lung metastasis, there was a clear decrease in the sh*Brd7* group (**SI 6H, J**) but not in the sh*Arid1A* group (**SI 6I, J**). We further confirmed this observed reduction in pulmonary tumor formation by depleting *Pbrm1* in the 4T1 isogenic cell line, 4T07. While these cells fail to metastasize from the fatpad, they do form robust pulmonary tumors upon tail vein injection^31^. Consistent with our findings in the 4T1 model, depletion of Pbrm1 in the 4T07 cells similarly reduce pulmonary tumor formation (**SI 6K, L**).

We next tested the effects of *Pbrm1* depletion in additional models of metastasis. We observed similar effects in metastasis from NME^30^ orthotopic tumors in NRG immunocompromised mice (**Fig 6K, L**) and found that metastases from the sg*Pbrm1* tumors had higher Pbrm1 expression relative to the parental line (**Fig 6M**), indicating that metastases enriched for populations of cells with residual Pbrm1 expression. We also observed a similar reduction in pulmonary tumors after tail vein injection with mouse syngeneic kidney cancer cell line Renca with *Pbrm1* deletion in the in BALB/c mice (**Fig 6N-Q; SI 6N-Q**) and in nu/nu mice (**SI 6R**), indicating that the reduction in metastasis in sg*Pbrm1* cells is not entirely dependent on immune function.

These findings strongly suggest that Pbrm1 facilitates metastasis *in vivo*, but do not provide clear insight into whether Pbrm1 is required for dissemination from the primary tumor, initiation, or outgrowth of pulmonary metastases. Therefore, we developed a 4T1 cell line with inducible expression of sh*Pbrm1* upon doxycycline administration (**Fig 6R, S**). These inducible sh*Pbrm1* cells were implanted into the mammary fat pads of Balb/c mice and primary tumors were allowed to grow for 2 weeks before surgical removal. After cellular dissemination and removal of the primary tumor, mice were administered doxycycline (**Fig 6T, U; SI 6S**). Consistent with our constitutive depletion models, doxycycline-induced depletion of Pbrm1 after primary tumor removal also reduced metastases (**Fig 6U**). This indicates that Pbrm1 required for the outgrowth of cells at the metastatic site.

### 13. Pbrm1-mediated gene expression in TNBC cells

To evaluate how the mechanistic insights regarding the genomic function of Pbrm1 in NMuMG cells undergoing EMT relate to the requirement for Pbrm1 in 4T1 metastasis, we performed RNA-seq on 4T1 cells treated with TGFβ1 (**SI 6T**). Although 4T1 cells are metastatic, they have a dynamic epithelial/mesenchymal phenotype that contributes to its aggressive nature and the ability to respond to TGFβ1. Like in NMuMG cells, all conditions segregated in the PCA plot (**Fig 6V).** Also similar to NMuMG cells, a large number of TGFβ1-inducible genes fail to be induced in the absence of *Pbrm1* (**Fig 6W**, highlighted, (**SI 6U**). Pathway analysis for this subset of genes enriched for processes such as cell proliferation, response to cytokines, migration, and extracellular matrix (**Fig 6X**), similar to NMuMG cells (**Fig 2K).** Further, in agreement with the findings in NMuMG cells, sh*Pbrm1* in untreated 4T1 cells increased invasive capabilities, while sh*Pbrm1* reduced invasive capabilities in TGFβ1 treated cells (**Fig 6Y**). Taken together, Pbrm1 is required for TGFβ1-dependent gene expression in both normal murine mammary cells and TNBC cells.

## Discussion

The mammalian PBAF subcomplex was biochemically defined at the same time as cBAF^20,94^, yet its biochemical role in gene transcription is much less clear. Here we find that in epithelial cells Pbrm1 deletion slightly increases the growth rate, but blocks completion of EMT upon stimulation with TGFβ1. This requirement for EMT is consistent with the phenotype described for the Pbrm1 knockout mouse, which is embryonic lethal at E14.5^95^ due to a failure in EMT during heart development^96,97^. It is also consistent with reports of a requirement for Pbrm1^98^ and ARID2^99^ in BMP/ TGFβ-dependent mesenchymal stem cell osteolineage differentiation and a requirement for Brd7 in the expression of TGFβ response genes^100^. Identifying this dependency on PBAF for EMT has allowed us the opportunity to dissect the role of PBAF in facilitating gene expression. We identified over 1,000 DEGs in epithelial cells with Pbrm1 knockout; however, the ChIP-Seq and ATAC-Seq provided very little insight into how PBAF might be involved in the regulation of these genes. As has been observed in other cell types^5,27^, PBAF is bound at promoters, with Phf10 binding correlating strongly with H3K4me3 enrichment. In NMuMG cells, we observed Phf10 binding at over 9,000 promoters; however, only about half of the DEGs have Phf10 enrichment at their promoters. Further, there was no difference of Phf10 enrichment in upregulated compared to downregulated genes, and no correlation between Phf10 enrichment and the degree to which gene expression changes upon Pbrm1 knockout. Upon treatment with TGFβ1, however, Phf10 is recruited to new sites, where we could detect significant changes in DNA accessibility using ATAC-Seq. Further, binding studies with recombinant nucleosomes reveal that, contrary to the results with histone peptides, only H3K14ac is important for Pbrm1 binding, which was supported by Phf10 enrichment at inducible sites marked only by H3K14ac. Although acetylation marks on histone 3 tails are often deposited indiscriminately by Gcn5^101^ and found together at promoters, H3K14ac is found alone at a subset of inducible genes^102,103^ indicating that PBAF activity may be more important for the initiation, rather than maintenance, of promoter accessibility. Uncovering the specific acetyltransferase responsible for depositing H3K14ac, and how its activated upon TGFβ1 treatment, will be useful for further establishing a role for PBAF in gene induction.

EMT is an important part of cancer metastasis, and we find that Pbrm1 dependency extends to the metastasis of 4T1 mouse breast cancer cells from the primary site to the lung. Using inducible *Pbrm1* knockdown, we establish that Pbrm1 is important for outgrowth of lung metastases after dissemination. These findings are consistent with our previous work demonstrating that cells must undergo an EMT prior to initiation of metastatic tumor growth^104^. What remains to be seen is whether deleting Pbrm1 in established metastatic tumors could halt or even regress metastatic tumor growth. *In vitro*, deletion of *Pbrm1* prevents the induction of a subset of TGFβ1 response genes in metastatic TNBC cells. In particular, the loss of Pbrm1 has a very pronounced effect on the expression of immunosuppressive^105,106^ IFNγ response genes during EMT. Pbrm1 regulation of IFNγ response genes was similarly observed in human metastatic TNBC cells^107^, metastatic prostate cancer cells^108^, and renal cancer cell lines^109–111^. This may implicate some contribution of the immune system in the reduction of metastases in *Pbrm1* depleted tumors. Consistent with this, PBAF subunit deletions in tumors increase T cell killing^112^ and predict a better response to immunotherapies^113,114^. This was originally hypothesized to be the result of an *increase* in interferon response in Pbrm1-mutant cells, either through Pbrm1-mediated gene repression^115^, Pbrm1-mediated DNA damage repair^116^, Pbrm1-mediated alleviation of replication stress^117,118^, or Pbrm1-mediated p53 regulation^119,120^; however, the expression of immunosuppressive genes may also contribute. Paradoxically, the loss of PBAF subunits can also result in immunologically cold tumors in some cases^121^, which is also attributed to the expression of IFNγ response genes^122^. Therefore the context-dependent roles for Pbrm1 in cancer progression may reflect the context-dependent roles for IFNγ ^123,124^ in cancer progression. Additional studies will be required to establish how Pbrm1 facilitates IFNγ response in tumors to facilitate metastasis and immune suppression, and the context in which pharmacologically targeting PBAF could be therapeutically advantageous.

## Limitations of this study

Transforming growth factor-β1 in NMuMGs induces EMT if cells are in G1/S, and induces apoptosis if cells are in G2/M^69^. Some of the phenotypes, and possibly gene expression changes, we observe with NMuMG cells may be related to the slightly increased proliferation rate or increased DNA-damage^125^ in the Pbrm1 knockout that increase apoptosis rates. Further genomic studies in the 4T1 cells, where we see no changes in cell cycle or apoptosis with Pbrm1 depletion, could give more insight into just the transcriptional roles for Pbrm1 during TGFβ1 stimulation. Further, the use of an inducible recruitment^7^ or inducible degron system would help separate direct from indirect gene targets.

While we observe a strong effect for Pbrm1 in breast cancer metastasis, this is likely context dependent. In renal cancers, Pbrm1 is mutated as an early event in the primary tumor after VHL mutation^126^. And while Pbrm1 mutant ccRCC tumors are much less metastatic^127^, they can eventually metastasize. In addition, recent reports find that Brd7 deletion in the 4T07 dormant breast cancer model permits the outgrowth of metastases^128^, implying that PBAF is not required for all metastatic tumors. Nevertheless, many reports indicate that Pbrm1 mutant tumors respond better to a variety of therapies^114,117,129^ implicating that targeting Pbrm1 as part of a combination therapy may be an effective treatment for metastatic disease. Further studies with pharmacological inhibitors of PBAF would resolve whether it is a potential therapeutic target in metastatic cancers as well as possibly in T cells to decrease exhaustion^130,131^.

## Acknowledgements

The authors thank past and current members of Dykhuizen lab and Wendt lab for their expertise and advice, Purdue Genomics Core, Purdue Cytometry and Cell Separation core and Purdue C3B core facilities for technical support. ECD is supported by NIH U01CA207532, V Foundation for Cancer Research (V2014-004 and D2016-030), American Cancer Society (RSG-21-012-01-DMC) and Indiana CTSI (TR001108). This work was supported by the Office of the Assistant Secretary of Defense for Health Affairs, through the Peer Reviewed Cancer Research Program, under Award No. W81XWH-17-1-0267 to ECD. Opinions, interpretations, conclusions, and recommendations are those of the author and are not necessarily endorsed by the Department of Defense. MKW is supported by NIH R01CA281216. VMW is supported by NIH R01EY033734. GJ is supported by a SIRG graduate fellowship from the Purdue Institute for Cancer Research. We acknowledge support from the IU Simon Cancer Center (Grant P30CA082709), Purdue University Center for Cancer Research (Grant P30CA023168) and Walther Cancer Foundation. *EpiCypher* is supported by the National Institutes of Health (NIH) through grants R44GM117683 and R44GM116584. Schematics were created using Biorender.

## Author Contributions

A.D., M.K.W. and E.C.D. conceived the experiments. A.D., M.G.A, J.L.M., A.B.S., W.A.T., V.M.W., M.R.M., M.K.W., and E.C.D. designed experiments. A.D., M.G.A, J.L.M., A.B.S., S.S.A., G.J., M.H., E.G.P., M.R.M., M.K.W., and E.C.D performed experiments and analysis. A.D., G.J., J.J-L., V.M.W., Y.Z., S.M.U., and E.C.D. performed bioinformatics analyses. M.H. and W.A.T. performed proteomics processing and analysis. A.D. and E.C.D. wrote the manuscript with assistance from M.K.W.

