## Supplemental Figures 1-6 for "PBRM1-Dependent PBAF Targeting is Required for EMT and Metastasis in Breast Cancer"

### Supplementary Figures

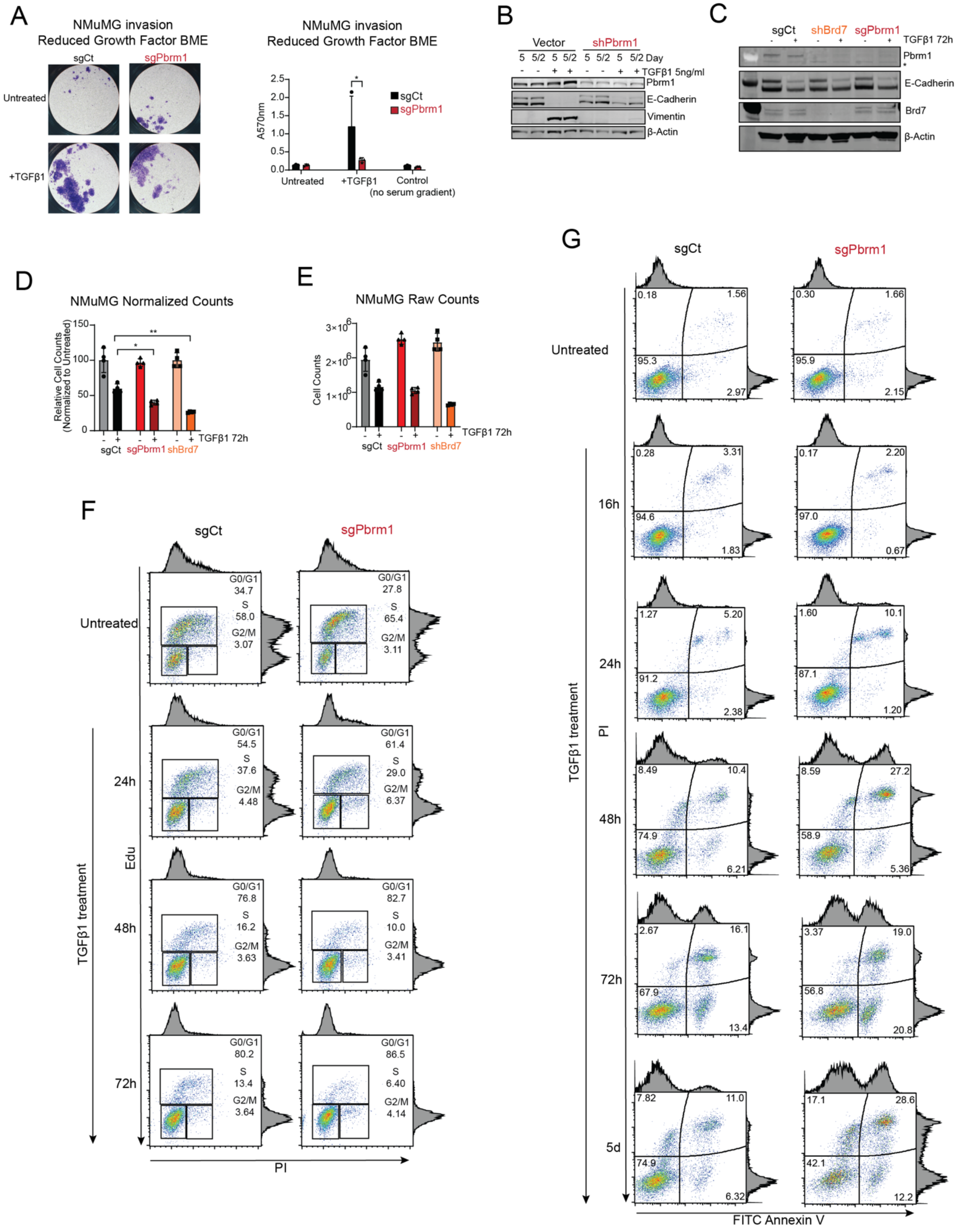

**SI Figure 1**

(A) Transwell invasion assay images and bar plot of absorbance quantification of NMuMG sgCt and sg*Pbrm1* cells with and without TGFβ1 treatment in reduced growth factor basement membrane (BME). n=3 technical replicates. Data are represented as mean ± SD.

(B) Immunoblots of Pbrm1, E-Cadherin, and Vimentin in NMuMG control and sh*Pbrm1* cells with and without TGFβ1 treatment with the indicated concentration and time periods. 5/2 denotes 5 days of TGFβ1 and next 2 days without TGFβ1.

(C) Immunoblots of Pbrm1 and E-Cadherin levels in NMuMG sgCt, sh*Brd7,* and sg*Pbrm1* cells with and without TGFβ1 treatment using whole cell extracts.

(D and E) Relative (D) and absolute (E) cell counts of NMuMG sgCt, sg*Pbrm1,* and sh*Brd7* cells with and without TGFβ1 treatment, normalized to untreated sample for relative counts. Representative graph, n=2 biological replicates. Data are represented as mean ± SD.

(F) Flow cytometry density dot plots of Edu-PI staining of percentage of cells in different cell cycle stages in NMuMG sgCt and sg*Pbrm1* cells with and without TGFβ1 treatment for the indicated time periods. n=3 biological replicates. Data are represented as mean ± SD.

(G) Flow cytometry density dot plots of AnnexinV-PI staining of percentage of live (AnnexinV- PI-, Q4), Annexin V+ (Q2 and Q3) and PI+ (Q1) cells in NMuMG sgCt and sg*Pbrm1* cells with and without TGFβ1 treatment for the indicated time periods. n=2 biological replicates. Data are represented as mean ± SD.

Statistical comparison was done using multiple unpaired t-tests with Holm-Sidak correction. ∗: $p < 0.05$, ∗∗: $p < 0.01$, ∗∗∗: $p < 0.001$, ∗∗∗∗: $p < 0.0001$

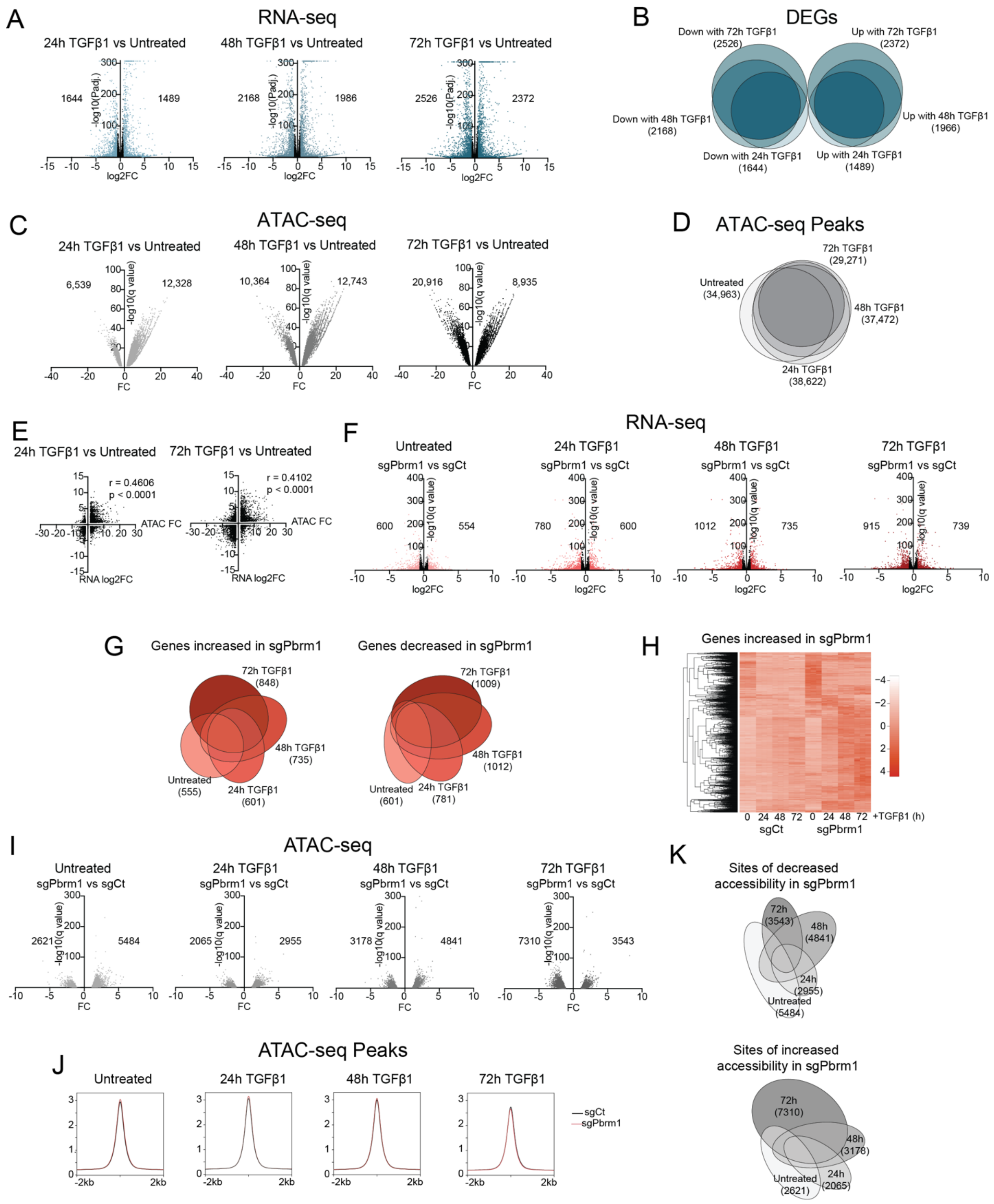


**SI Figure 2**

(A) Volcano plots of DEGs (padj<0.05) identified in NMuMG sgCt cells with TGFβ1 treatment between the indicated comparisons using RNA-seq. DEGs with |log2FC|>1.5 are shown in gradient of blue, each corresponding to a specific TGFβ1 treatment time.

(B) Venn diagram of DEGs shared between the indicated conditions, as identified in RNA-seq. Total number of DEGs in each condition is indicated in parentheses.

(C) Volcano plots of number of regions displaying differential accessibility identified in NMuMG sgCt cells with TGFβ1 treatment between the indicated comparisons using ATAC-seq. Regions with differential accessibility (padj<0.05, |FC|>1.5) are shown in gradient of black, each corresponding to a specific comparison.

(D) Venn diagram of accessible regions, as identified in ATAC-seq. Total number of regions in each condition is indicated in parentheses.

(E) Scatter plot of differentially accessible regions from ATAC-seq with differential expression of the nearest gene from RNA-Seq, at 24h (left) and 72h (right) of TGFβ1 treatment compared to untreated NMuMG sgCt cells. Each data point in the scatter plot represents on the *x axis*: differentially accessible regions as FC from ATAC-seq and on the *y axis*: expression change of the nearest gene from RNA-seq as log2FC with 24 or 72h TGFβ1 treatment relative to no treatment.

(F) Volcano plots of DEGs (padj<0.05) identified in NMuMG sg*Pbrm1* relative to sgCt cells in untreated, 24, 48, and 72h TGFβ1 treated conditions using RNA-seq. DEGs with |log2FC|>1.5 are shown in shades of red, with the number of genes increased and decreased in expression indicated in the plot.

(G) Venn diagram of DEGs shared between the indicated conditions, as identified in RNA-seq. Total number of DEGs in each condition is indicated in parentheses.

(H) Heatmap representation of all genes increased in NMuMG sg*Pbrm1* relative to sgCt cells at any timepoint of TGFβ1 treatment compared to untreated cells.

(I) Volcano plots of differentially accessible regions identified in NMuMG sg*Pbrm1* relative to sgCt cells in untreated, 24, 48, and 72h TGFβ1 treated conditions using ATAC-seq. The number of regions with significantly increased and decreased accessibility is indicated in the plot.

(J) Metagene plots of global accessibility in NMuMG sgCt and sg*Pbrm1* cells in untreated and TGFβ1 treated conditions as identified in ATAC-seq. The regions used for the average peak size are the overlap of all peaks identified from any condition/genotype

(K) Venn diagram of differentially accessible regions (decreased (top) or increased (bottom)) shared between the indicated conditions, as identified in ATAC-seq. Total number of regions in each condition is indicated in parentheses.

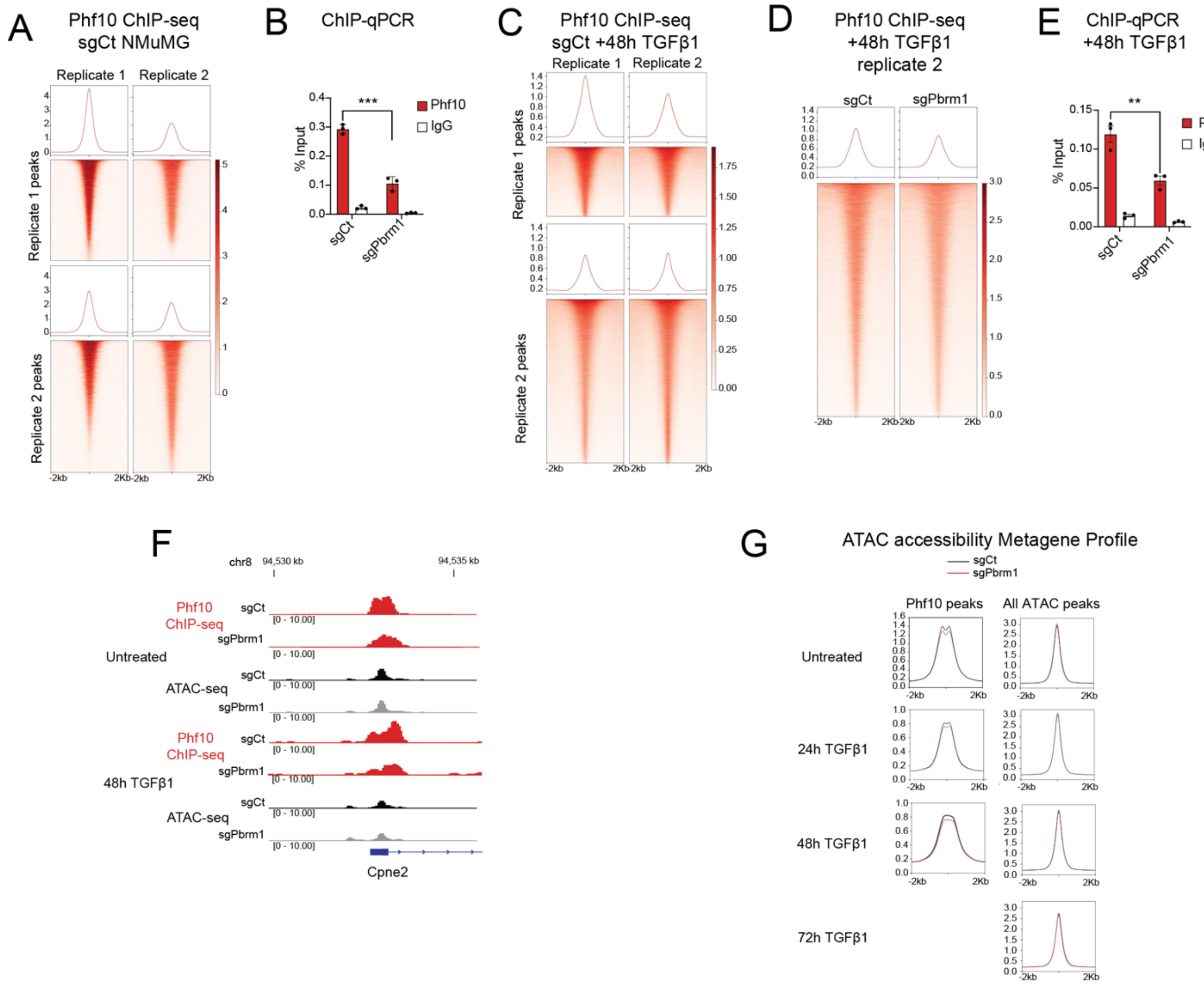


**SI Figure 3**

(A) Metagene plots and heatmaps of ChIP-seq enrichment of Phf10 at Phf10 sites identified in two independent ChIP-seq replicates in untreated NMuMG sgCt cells.

(B) Bar plot of the percent of input DNA enriched from ChIP of Phf10 and IgG in NMuMG sgCt and sg*Pbrm1* cells using qPCR. N = 3 independent replicates.

(C) Metagene plots and heatmaps of ChIP-seq enrichment of Phf10 at Phf10 sites identified in two independent ChIP-seq replicates in 48h TGFβ1 treated NMuMG sgCt cells.

(D) Metagene plots and heatmaps of ChIP-seq enrichment of Phf10 in 48h TGFβ1 treated NMuMG sgCt and sg*Pbrm1* cells using Phf10 peaks identified in NMuMG sgCt cells with 48h TGFβ1.

(E) Bar plot of the percent of input DNA enriched from ChIP of Phf10 and IgG in NMuMG sgCt and sg*Pbrm1* cells with 48h TGFβ1 treatment using qPCR. N = 3 independent replicates.

(F) Genomic tracks of ChIP-seq enrichment of Phf10 and ATAC-seq profile in untreated and 48h TGFβ1-treated NMuMG sgCt and sg*Pbrm1* cells at *Cpne2* locus

(G) Metagene plots of accessibility profile of NMuMG sgCt and sg*Pbrm1* cells with 0-72h TGFβ1 treatment at all regions identified in ATAC-seq, compared to accessibility profile at Phf10 ChIP-seq peaks identified in NMuMG sgCt cells with 0, 24 or 48h TGFβ1 treatment.

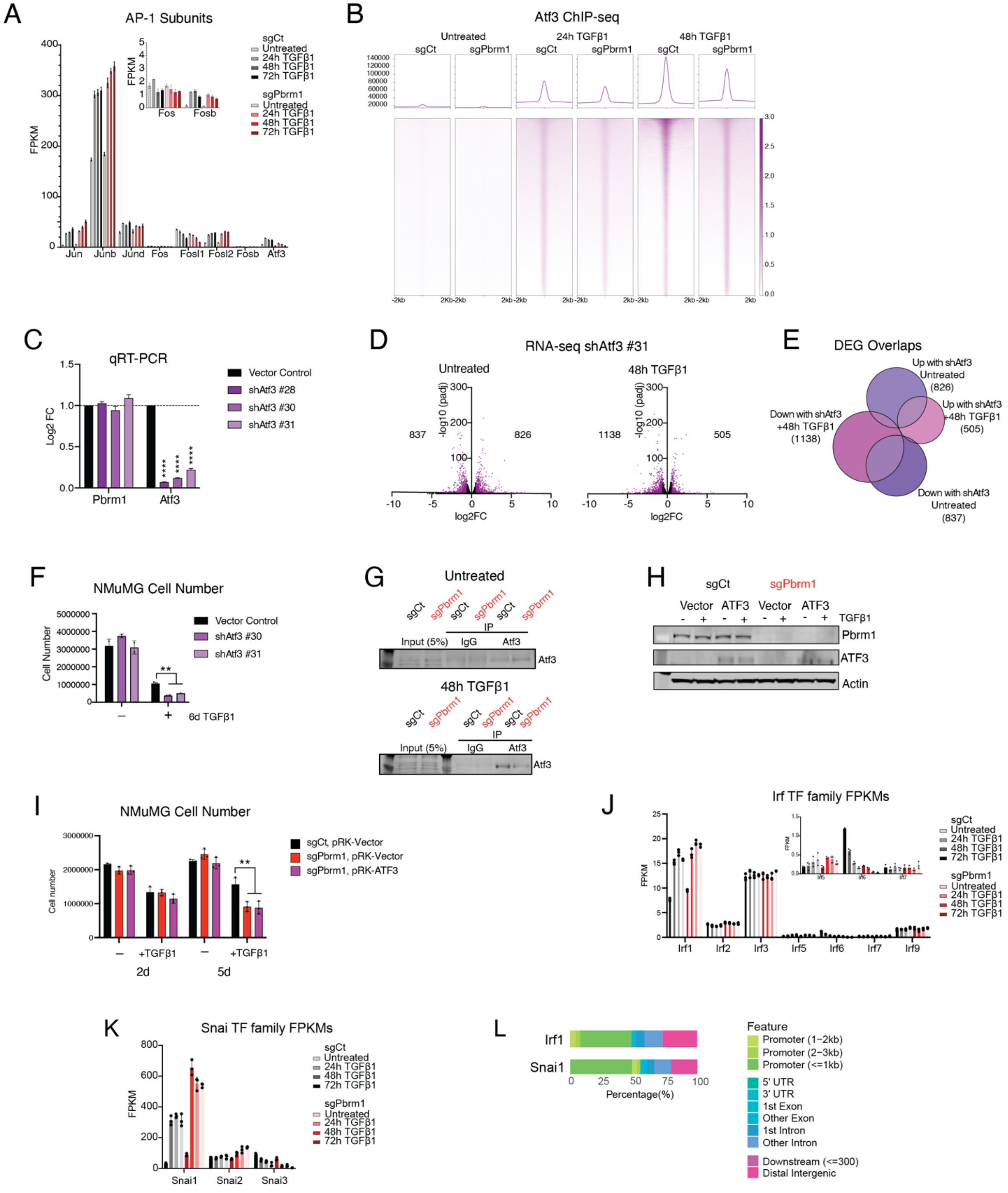


**SI Figure 4**

(A) Bar plot of FPKM values for the different AP-1 subunits from RNA-seq of NMuMG sgCt and sg*Pbrm1* cells with 0, 24, 48, and 72h TGFβ1 treatment.

(B) Metagene plots and heatmaps of ChIP-seq enrichment of Atf3 in NMuMG sgCt and sg*Pbrm1* cells with 0, 24, and 48h TGFβ1 treatment.

(C) qRT-PCR of NMuMG cells treated with three different shRNA against *Atf3*. *Oaz1* was used as the housekeeping gene. Data are represented as mean ± SD.

(D) Volcano plots of DEGs (padj.<0.05) identified in NMuMG sh*Atf3* relative to control cells in untreated and 48h TGFβ1 treated conditions using RNA-seq. DEGs with |log2FC|>1.5 are shown in shades of magenta, with the number of genes increased and decreased in expression indicated in the plot.

(E) Venn diagram of DEGs from RNA-seq of sh*Atf3* relative to control cells in both untreated and 48h TGFβ1-treated cells. Total number of DEGs in each condition is indicated in parentheses.

(F) Absolute cell counts of NMuMG control and sh*Atf3* cells with and without TGFβ1 treatment. 0.9 million cells/well were seeded for each cell line and cell counts were taken on day 6. Representative graph, n=3 biological replicates. Data are represented as mean ± SD.

(G) Immunoblots of lysates and immunoprecipitations from NMuMG sgCt and sg*Pbrm1* cells in untreated and 48h TGFβ1 treated conditions. n=2 biological replicates.

(H) Immunoblots of whole cell extracts from NMuMG sgCt and sg*Pbrm1* cells with human ATF3 expression.

(I) Absolute cell counts of NMuMG sgCt, sg*Pbrm1* and sg*Pbrm1*-*ATF3* re-expressing cells with and without 5 ng/mL TGFβ1 treatment. 0.8 million cells/well were seeded for each cell line and cell counts were taken on day 2 and 5. Representative graph, n=4 biological replicates. Data are represented as mean ± SD.

(J and K) Bar plots of FPKM values for the Irf (D) and Snai (E) TFs from RNA-seq of NMuMG sgCt and sg*Pbrm1* cells with 0, 24, 48, and 72h TGFβ1 treatment.

(L) Genomic feature distribution of the ChIP-seq peaks identified for Irf1 in NMuMG cells with 48h TGFβ1, and Snai1 in pBI.3G cells.

Statistical comparison was done using multiple unpaired t-tests with Holm-Sidak correction. ∗: p < 0.05, ∗∗: p < 0.01, ∗∗∗: p < 0.001, ∗∗∗∗: p < 0.0001

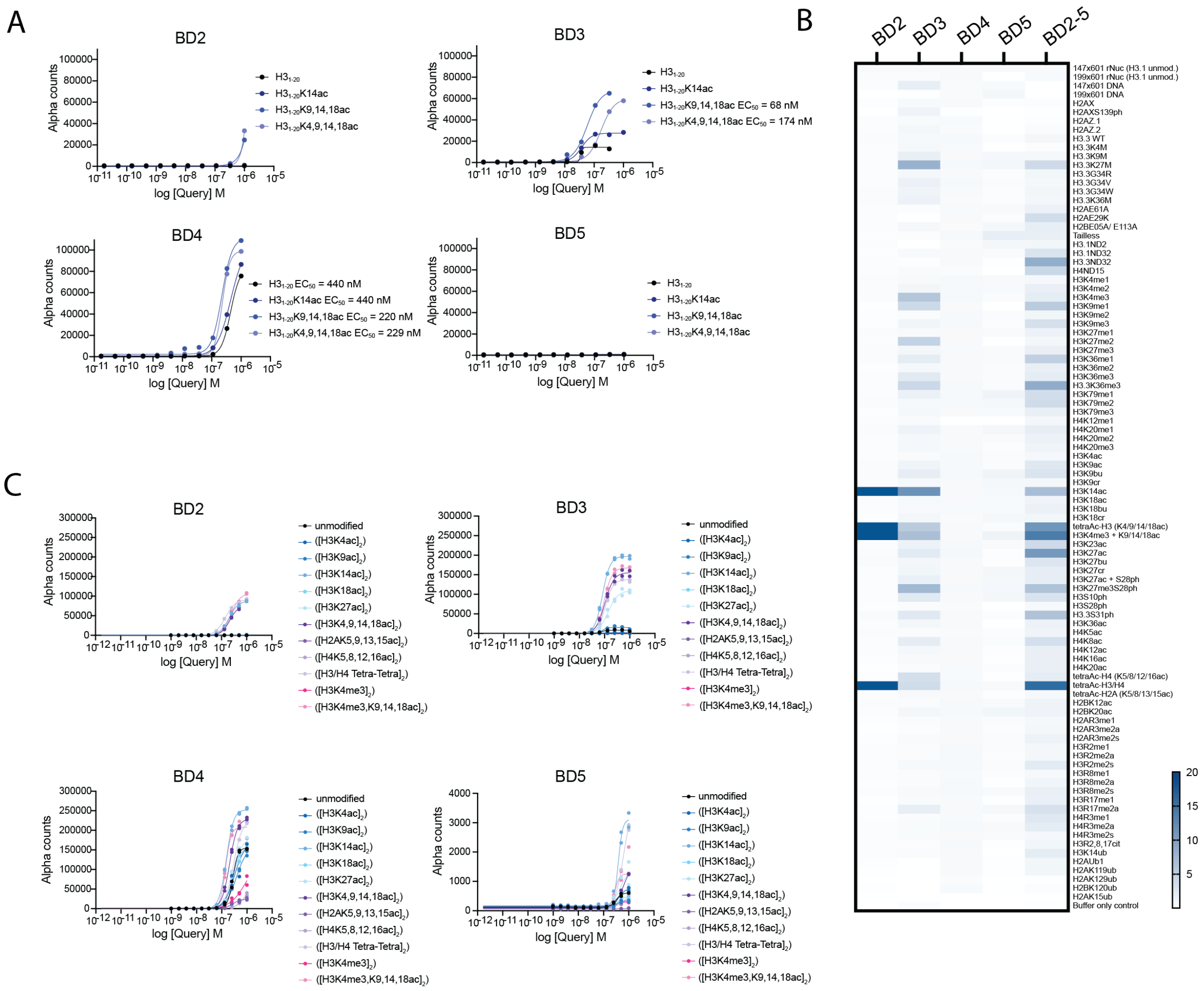


### SI Figure 5

(A) Binding curves of BD2, BD3, BD4, and BD5 with the indicated peptides obtained using the Captify™ assay. $EC_{50}$ (nM) values are indicated next to the corresponding curves.

(B) Heatmap representation of signal in the Captify™ assay with BD2, BD3, BD4, BD5, and tandem BD2-5 and nucleosomes with the indicated modifications. The data presented as Alpha Counts normalized to the signal from unmodified 147x601 rNucs.

(C) Binding curves of BD2, BD3, BD4 and BD5 for nucleosomes bearing the indicated peptides obtained using the Captify™ assay. $EC_{50}$ (nM) values are listed in the table in Fig 5E.

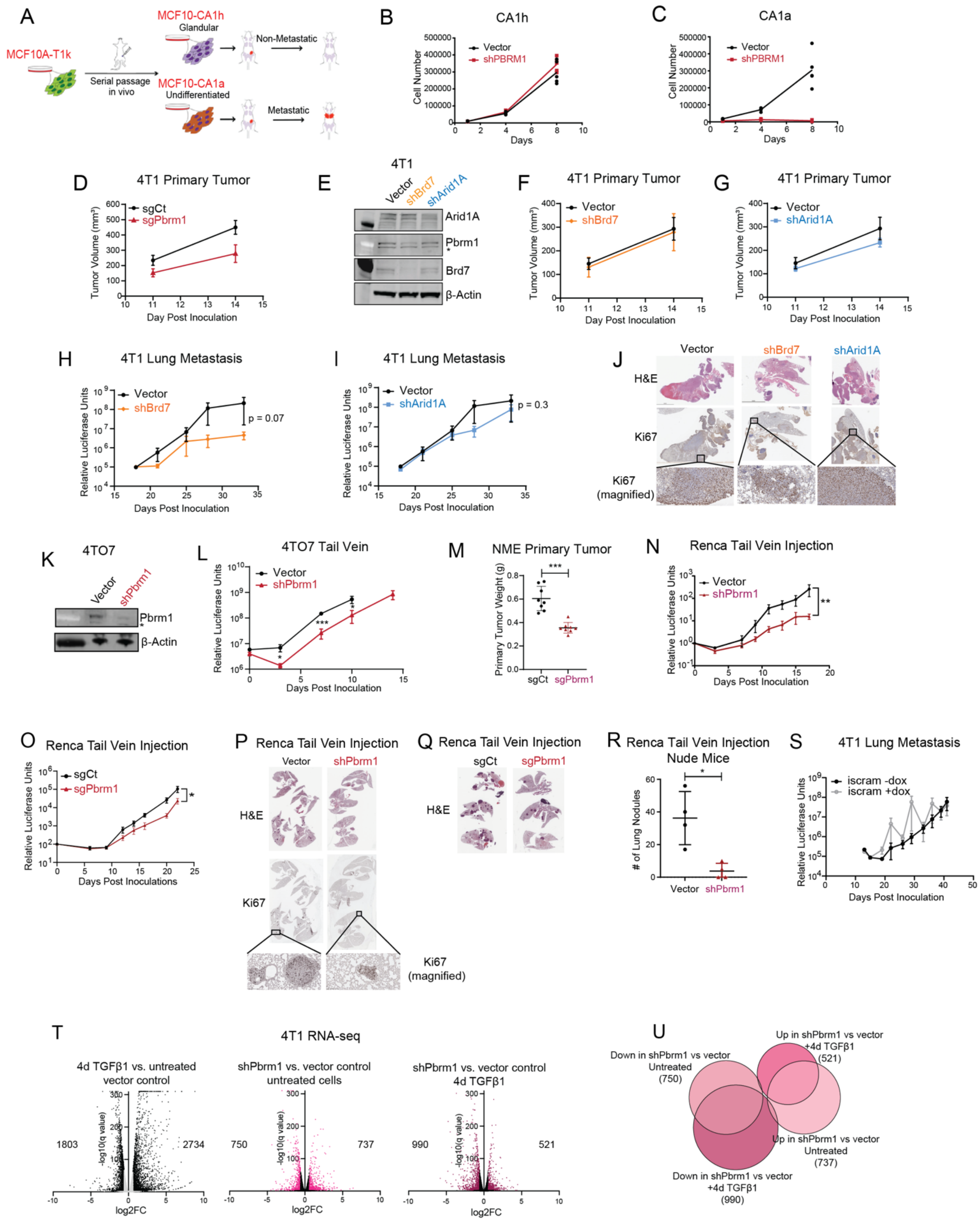

**SI Figure 6**

(A) Schematic diagram of the development of MCF10 series of cell lines- T1K, CA1h, and CA1a.

(B and C) *In vitro* proliferation of MCF10 CA1h (B) and CA1a (C) control and sh*Pbrm1* cells; 10,000 cells were plated at day 0. n=3 biological replicates. Two-way ANOVA was used for statistical comparison. Data are represented as mean ± SD.

(D) Primary tumor growth in 4T1 sgCt and sg*Pbrm1* cells measured using vernier calipers at the indicated time points. Two-way ANOVA with multiple comparisons was used for statistical comparison. Data are represented as mean ± SEM.

(E) Immunoblots of whole cell lysates from 4T1 cells expressing control vector, sh*Brd7* and sh*Arid1A*.

(F and G) Primary tumor growth in 4T1 sh*Brd7* (F) and sh*Arid1A* (G) relative to control vector cells, measured using vernier calipers at the indicated time points. Two-way ANOVA with multiple comparisons was used for statistical comparison. Data are represented as mean ± SEM.

(H and I) Bioluminescent imaging of lung metastasis in 4T1 sh*Brd7* (H) and sh*Arid1A* (I) relative to vector control cells after removal of the primary tumor. Two-way ANOVA with multiple comparisons was used for statistical comparison. Data are represented as mean ± SEM.

(J) H&E and Ki67 stained IHC images of the lungs harvested at the end of the experiment described in (H) and (I).

(K) Immunoblots of whole cell lysates from 4TO7 cells expressing sh*Pbrm1* or vector control.

(L) Bioluminescent imaging of lung metastasis in 4TO7 vector control and sh*Pbrm1* cells. Welch's t-test was used for statistical comparison. Data are represented as mean ± SEM.

(M) Scatter plot of primary tumor weights from individual mice in the NME sgCt and sg*Pbrm1* groups harvested at the end of experiment. Welch's t-test was used for statistical comparison. Data are represented as mean ± SD.

(N and O) Bioluminescent imaging of lung metastasis in Renca sh*Pbrm1* (N) and sg*Pbrm1* (O) relative to control cells. Two-way ANOVA with multiple comparisons was used for statistical comparison. Data are represented as mean ± SEM.

(P) H&E and Ki67 stained IHC images of the lungs harvested at the end of the experiment described in (N).

(Q) H&E images of the lungs harvested at the end of the experiment described in (O).

(R) Scatter plot of number of lung nodules from individual mice in the Renca control and sh*Pbrm1* groups harvested at the end of experiment. Welch's t-test was used for statistical comparison. Data are represented as mean ± SD.

(S) Bioluminescent imaging of lung metastasis in 4T1 iscram cells with and without doxycycline administration, after removal of the primary tumor. Two-way ANOVA with multiple comparisons was used for statistical comparison. Data are represented as mean ± SEM.

(T) Volcano plots of DEGs (padj.<0.05) identified from RNA-Seq of 4T1 vector control cells with 4d TGFβ1 treatment compared to untreated cells (left), sh*Pbrm1* relative to vector control cells in untreated conditions (middle), and sh*Pbrm1* relative to vector control cells with 4d TGFβ1 (right). DEGs with |log2FC|>1.5 are shown in color with the number of genes increased and decreased in expression indicated in the plot.

(U) Venn diagram of DEGs identified in RNA-seq of 4T1 cells with untreated sh*Pbrm1* relative to vector control cells and 4d TGFβ1 treated sh*Pbrm1* relative to vector control cells. Total number of DEGs in each condition is indicated in parentheses.

∗: $p < 0.05$, ∗∗: $p < 0.01$, ∗∗∗: $p < 0.001$, ∗∗∗∗: $p < 0.0001$
